# Interhemispheric CA1 projections support spatial cognition and are affected in a mouse model of the 22q11.2 deletion syndrome

**DOI:** 10.1101/2024.09.05.611389

**Authors:** Noelia S. de León Reyes, Maria Helena Bortolozzo-Gleich, Yuki Nomura, Cristina García Fregola, Marta Nieto, Joseph A. Gogos, Félix Leroy

## Abstract

Untangling the hippocampus connectivity is critical for understanding the mechanisms supporting learning and memory. However, the function of interhemispheric connections between hippocampal formations is still poorly understood. So far, two major hippocampal commissural projections have been characterized in rodents. Mossy cells from the hilus of the dentate gyrus project to the inner molecular layer of the contralateral dentate gyrus and CA3 and CA2 pyramidal neuron axonal collaterals to contralateral CA3, CA2 and CA1. In contrary, little is known about commissural projection from the CA1 region. Here, we show that CA1 pyramidal neurons from the dorsal hippocampus project to contralateral dorsal CA1 as well as dorsal subiculum. We further demonstrate that the interhemispheric projection from CA1 to dorsal subiculum supports spatial memory and spatial working memory in WT mice, two cognitive functions impaired in male mice from the *Df16(A)^+/-^* model of 22q11.2 deletion syndrome (22q11.2DS) associated with schizophrenia. Investigation of the CA1 interhemispheric projections in *Df16(A)^+/-^* mice revealed that these projections are disrupted with male mutants showing stronger anatomical defects compared to females. Overall, our results characterize a novel interhemispheric projection from dCA1 to dorsal subiculum and suggest that dysregulation of this projection may contribute to the cognitive deficits associated with the 22q11.2DS.

## Introduction

Exploring brain circuitry is a major endeavor of neuroscience but most studies focus on connections between regions located on one side of the brain. Interhemispheric connections are however essentials in bilateral animals, including in vertebrates.^1^ Bilateral integration allows computation of information from the two hemispheres and results in a more complex output than the one provided by individual inputs from each hemisphere.^2^ Furthermore, an increase in commissural projections accompanies the evolutionary increase in brain size and connectivity,^3^ probably to help synchronize neuronal activity between brain hemispheres. Indeed, as bilateral brain regions support slightly different functions, a phenomenon called lateralization,^4–6^ interhemispheric connections are also key to coordinate neuronal activity. Accordingly, dysfunction in the transfer of information between the two cerebral hemispheres has been implicated in a number of neurodevelopmental and psychiatric disorders.^7–10^

The hippocampus, a bilateral structure critical for episodic memory exhibits asymmetry at molecular and functional levels.^11^ Thus, silencing CA3 in the left hemisphere hippocampus of mice is sufficient to impair spatial long-term memory but silencing the contralateral CA3 in the right hemisphere has no effect.^12^ In humans, the amplitude of low-theta oscillations increases in the left but not in the right hippocampus when remembering object–location pairs.^13^ In contrast, low-theta activity increases in the right but not in the left hippocampus during periods of navigation.^13^ Overall, hippocampal lateralization highlights the importance of understanding how left and right hippocampus formation process information differently and communicate with each other.

Dorsal hippocampi are massively interconnected through two interhemispheric pathways: the ventral and dorsal hippocampal commissures,^14^ whose names refer to their dorsoventral location with respect to the commissural plate during development.^15^ In adult rodents, the ventral hippocampal commissure is located at the anterior part of the fornix and harbors most of the interhemispheric connections between hippocampi. The dorsal hippocampal commissure on the other hand is located more posterior, closer to the splenium of the corpus callosum and contains mostly fibers originating from the contralateral parahippocampus.^16^ Interhemispheric hippocampal projections are critical to several hippocampal functions such as recognition,^17^ contextual and spatial memories^18^ as well as generalization.^19^ For example, monkeys whose dorsal hippocampal commissure was sectioned made more errors and showed evident learning difficulties in a visual discrimination task.^20^ More recently, it has been shown that individual variations in human and non-human primate dorsal commissure correlate with performance in a standardized recognition task.^21^ At the cellular level, mossy cells from the hilus of the dentate gyrus form connections with granule cells of the molecular layer located in the contralateral dentate gyrus.^22^ In addition, contralateral dentate gyrus also receives weak and sparse inputs from somatostatin-expressing inhibitory neurons located in the ipsilateral dentate gyrus^23,24^ and manipulating this projection disrupts contextual and spatial memories.^25^ The axons of CA3 and CA2 pyramidal neurons branch extensively, sending several collaterals toward the contralateral hippocampus.^26,27^ The function of these collaterals remains understudied, but they are believed to support synchronization of activity and the pattern completion property associated with dorsal CA3.^28^ Silencing contralateral projections from the right CA3 to the left CA1 show that this projection is necessary for long-term memory formation,^29^ which reinforces the idea that the left hippocampus has a predominant role in long-term memory formation.^12^ Moreover, slice physiology demonstrates that synapses from the left CA3 pyramidal neurons to the right CA1 pyramidal neurons exhibit plasticity unlike synapses from the right CA3 pyramidal neurons to the left CA1 pyramidal neurons.^30^ Despite evidence of asymmetry, the functional consequence of CA3 lateralization remains unknown. Finally, dorsal CA1 (dCA1) is essential for episodic memory^31^ and generalization^32,33^ and a previous investigation reported that some dCA1 pyramidal neurons project to contralateral dCA1 to govern rapid generalization but not fear memory (a form of episodic memory).^19^ This suggests different functions for ipsi- and contralateral projections originating from dCA1.

Adults and children with the 22q11.2DS demonstrate an array of cognitive deficits^34,35^ and a marked, 30-fold increase in the risk of developing schizophrenia during adolescence and early adulthood.^36,37^ Children with 22q11.2DS also exhibit and increased prevalence of attention-deficit hyperactivity disorder, autism spectrum disorder, mood and anxiety disorders, seizures and epilepsy.^38–40^ Cognitive dysfunction, a key manifestation of SCZ, is highly correlated with functional outcome and is a robust indicator of the risk of developing a psychotic illness.^41,42^ The hippocampus supports cognitive functions such as working or episodic memory which are impaired in SCZ.^43^ In addition, postmortem and in vivo neuroimaging studies in human have described an early involvement of the hippocampus in the pathophysiology of SCZ and suggested that dysregulation of glutamate neurotransmission originating in the hippocampal CA1 region may spread to downstream regions and initiate the transition from attenuated to syndromal psychosis.^44^ Brain imaging studies of 22q11DS patients also reported alterations in the anterior hippocampus,^45^ disrupted fornix integrity^46^ and developmental dysconnectivity.^47^ Accordingly, many mouse models of SCZ etiology, including the *Df16(A)^+/-^* mouse model of the 22q11.2DS, which carries a hemizygous 1.3 Mb deficiency that simulates the 1.5 Mb human microdeletion),^48^ exhibit hippocampal alterations.^49–51^ Thus, CA1 pyramidal neurons of *Df16(A)^+/-^* mice show changes in their dendritic tree, spine maturity, electrophysiological properties and receive less inhibitory inputs.^47,48,52^ Consequently, CA1 interneurons carry markedly reduced spatial information during random foraging^53^ and CA1 place cell dynamics are impaired.^54^ At behavioral level, *Df16(A)^+/-^* mice exhibit behavioral deficits in hippocampal-related behaviors such as fear conditioning,^48^ spatial working memory^55^ and social memory.^56^ Despite the volume of work, the role that hippocampal commissural projections play in the emergence of behavioral phenotypes exhibited by mouse models of SCZ etiology, including the *Df16(A)^+/-^* mouse model, remains unknown.

Here, we show that ipsilateral dCA1 pyramidal neurons project to contralateral CA1 and contralateral dorsal subiculum. We further demonstrate that silencing the commissural projections from dCA1 to dorsal subiculum impairs spatial memory and spatial working memory. As altered spatial cognition is a hallmark of 22q11.2DS, we characterized the performance of *Df16(A)^+/-^* mice from both sexes and found male mutants to be preferentially impaired. Finally, anatomical investigation revealed that interhemispheric projections are disrupted with male mutants showing stronger anatomical defects compared to females. Overall, our results characterize a novel interhemispheric projection from dCA1 pyramidal neurons to dorsal subiculum which supports spatial cognition and is affected by a mutation predisposing to cognitive dysfunction.

## Results

### dCA1 pyramidal neurons project to contralateral dCA1 and contralateral dorsal subiculum

In order to trace the outputs of CA1 pyramidal neurons, we injected the right dCA1 of *Lypd1-Cre* mice^57^ with a Cre-dependent AAV expressing membrane-bound GFP to label axonal fibers and synaptophysin tagged with mRuby to label synaptic terminals (Fig. 1a-b). We used the Satb2 marker of excitatory neurons^58^ to confirm that Cre expression is restricted to CA1 pyramidal neurons in the hippocampus proper. Indeed, when injecting a Cre-dependent virus expressing GFP, no recombination was observed in dCA2, dCA3 or in dCA1 GABAergic cells (Fig. S1), which is consistent with previous characterization of the mouse line.^59^ GFP expression was particularly prominent in pyramidal neurons from the deep layer of dCA1 stratum pyramidale (Fig. 1c-e). We then characterized the interhemispheric projections in the contralateral hippocampus (Fig. 1f-n). In the septal pole, we observed fibers and synaptic terminals in dCA1 (Fig. 1g-h)^19^. The majority of these terminals were located in the stratum oriens (s.o.) with only very sparse innervation in the stratum radiatum (s.r, Fig. 1g-k). In more temporal sections however, mRuby^+^ synaptic terminals were exclusively present in the dorsal subiculum (Fig. 1m-n). In the ipsilateral hippocampus, we saw abundant projections into dorsal subiculum and in layer V of the entorhinal cortex (Fig. S2). Outside the hippocampal formation, we also observed contralateral interhemispheric projections into several midline nuclei such as the lateral septum, nucleus reuniens, rhomboid nucleus and the latero-dorsal nucleus of the thalamus (Fig. S3). However, we did not observe terminals in the contralateral entorhinal cortex (Fig. S2). Upon close inspection of the ventral and dorsal hippocampal commissures, we detected dCA1 axons crossing the midline through both of them (Fig. 1g-j & S4a-d), suggesting that dCA1 interhemispheric neurons might use one route or the other according to the location of their contralateral targets. Injections targeting ventral CA1 (vCA1) in the temporal pole of the hippocampus showed no projections to the contralateral hippocampal formation (Fig. S4e-g). Overall, these experiments show that dCA1 and dorsal subiculum are the main targets of dCA1 interhemispheric projections.

**Figure 1.**
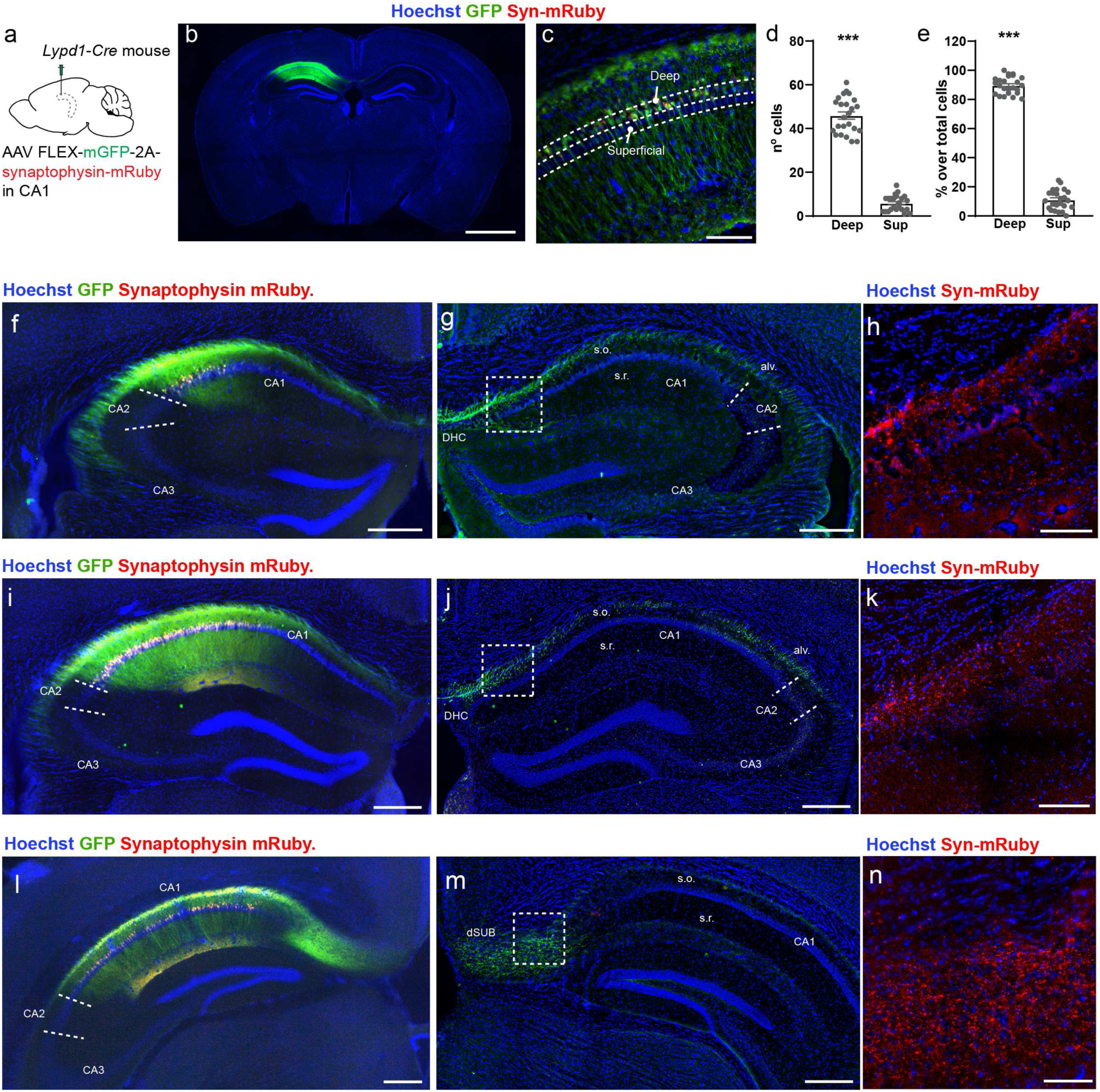
dCA1 pyramidal neurons project to contralateral dorsal CA1 and subiculum. **a.** *Lypd1-Cre* mice injected in dCA1 with AAV2/DJ hSyn.FLEX.mGFP.2A.Synatophysin-mRuby. **b-c.** Coronal section labelled for GFP and mRuby. **d-e.** Number and percentage of GFP^+^ cells in deep and superficial layers of dCA1. Each point corresponds to one observation (4 mice, 3 sections per mouse). **f-n.** Coronal hippocampal sections showing GFP^+^ cells in anterior (f), medial (i) and posterior ipsilateral dCA1 (l) and GFP^+^ fibers and mRuby^+^ pre-synaptic terminals in contralateral dCA1 and dSUB. For the entire figure, bar graphs represent mean ± SEM. Scale bars: 1mm (b), 300 µm (f,g,i,j,l,m) and 100 µm (c,h,k,n). (alv.): alveus, (s.o): stratum oriens and (s.r.): stratum radiatum.

While the emergence and temporal dynamics of the corpus callosum development are well-characterized,^60^ little is known about the development of the hippocampal interhemispheric projections. We performed in utero electroporation of a plasmid expressing GFP in the right lumen of wild-type mice at E14 and observed the brains at P7 in order to visualize interhemispheric hippocampal axons at a representative mid-developmental stage (Fig. S5). We took advantage of the third-electrode probe system^61^ to electroporate either dCA1 or the entire hippocampus to compare interhemispheric projections originating from dCA1 only vs. projections from CA1-CA3 and DG. In the first configuration of electroporation (Fig. S5a), the majority of GFP^+^ cells were located in dCA1 (Fig. S5b) and sent many axons through the dorsal hippocampus commissure which reached the stratum oriens of contralateral dCA1 (Fig. S5b-c). No axons were observed in the contralateral DG (Fig. S5d). Some axons coursed the ventral hippocampal commissure too (Fig. S5e) suggesting that at P7 the innervation pattern is already similar to what we observed in the adult (Fig. 1g). In the second configuration (Fig. S5f) we labeled the entire hippocampus (Fig. S5g). In this case, we observed GFP^+^ fibers in the stratum radiatum and stratum oriens of the entire contralateral hippocampal formation (Fig. S5g-h), as well as in the contralateral DG (Fig. S5i) which is consistent with the innervation pattern of Schaffer collaterals originating from dCA3 and mossy cells from DG (targeting the contralateral hilus). Compared to our CA1-specific targeting, we observed a higher number of fibers crossing the midline through the ventral hippocampal commissure (Fig. S5j). We concluded that interhemispheric projections from dCA1 emerge early in development and reach their final targets already at P7, with the majority of the projections crossing the midline through the dorsal hippocampal commissure, while the main crossing route for dCA3 and dCA2 interhemispheric axons is the ventral commissure.

### Interhemispheric projections from dCA1 to dorsal subiculum support spatial memory

Zhou et al.^19^ previously investigated the function of the interhemispheric dCA1 projection to dCA1 but the function of the interhemispheric dCA1 projection to subiculum is unknown. As dorsal CA1 and subiculum are both critical for spatial memory and spatial working memory,^62,63^ we tested whether silencing the interhemispheric dCA1 projection to subiculum would impair these cognitive functions. To this end, we performed the object location test of spatial memory^64^ and the spontaneous alternation T-maze test for spatial working memory^65^ on *Lypd1-*Cre mice. We also performed the novel object recognition test^66^ as a non-spatial memory control. We used a novel silencing opsin targeted to synaptic terminal^67^ to silence the dCA1 projection to contralateral dSUB going from the right hemisphere to the left one. Specifically, we injected the right dCA1 of *Lypd1-Cre* or WT mice with a Cre-dependent virus expressing the mosquito opsin eOPN3 tagged with mScarlet (Fig. 2a). As expected, only *Lypd1-Cre* mice showed viral expression (Fig. S6). For these experiments, we only included the mice with optic fiber implants above the left dorsal subiculum (Fig. S6). When using, light was applied during all trials. *Lypd1*-Cre mice performed the object location test, which consists of habituating the mouse to the same object in several locations of an open field (learning phase: 1^st^ to 3^rd^ trial, Fig. 2b).^64^ During the test phase (4^th^ trial), the mice have the option to explore the objects in a familiar or novel location. In control groups, mice preferred to interact with the object in the novel location (Fig. 2c-d), indicating normal spatial memory. Mice for which the dCA1 projection to contralateral dSUB was silenced *(Lypd1-Cre* with light) interacted to the same extent with objects in novel and familiar locations and did not exhibit any preference (Fig. 2c-d). Furthermore, *Lypd1-Cre* but not WT mice exhibited a decrease in discrimination index in light-on condition compared to the light-off condition (Fig. 2e-f). Silencing the projection therefore impaired the performance of the mice in this test of spatial memory. This was not due to changes in locomotion or interaction time with objects during trial 4 (Fig. 2g-h). In addition, the distance traveled decreased similarly across habituation trials, which indicates normal habituation to the arena (Fig. 2i).^68^ However, *Lypd1-Cre* mice in light-on condition exhibited an increase in interaction time during the repetitive presentation of the object which was not seen in control groups (Fig. 2j). Overall, this experiment shows that silencing interhemispheric projections from dCA1 to the dorsal subiculum impairs spatial memory.

**Figure 2.**
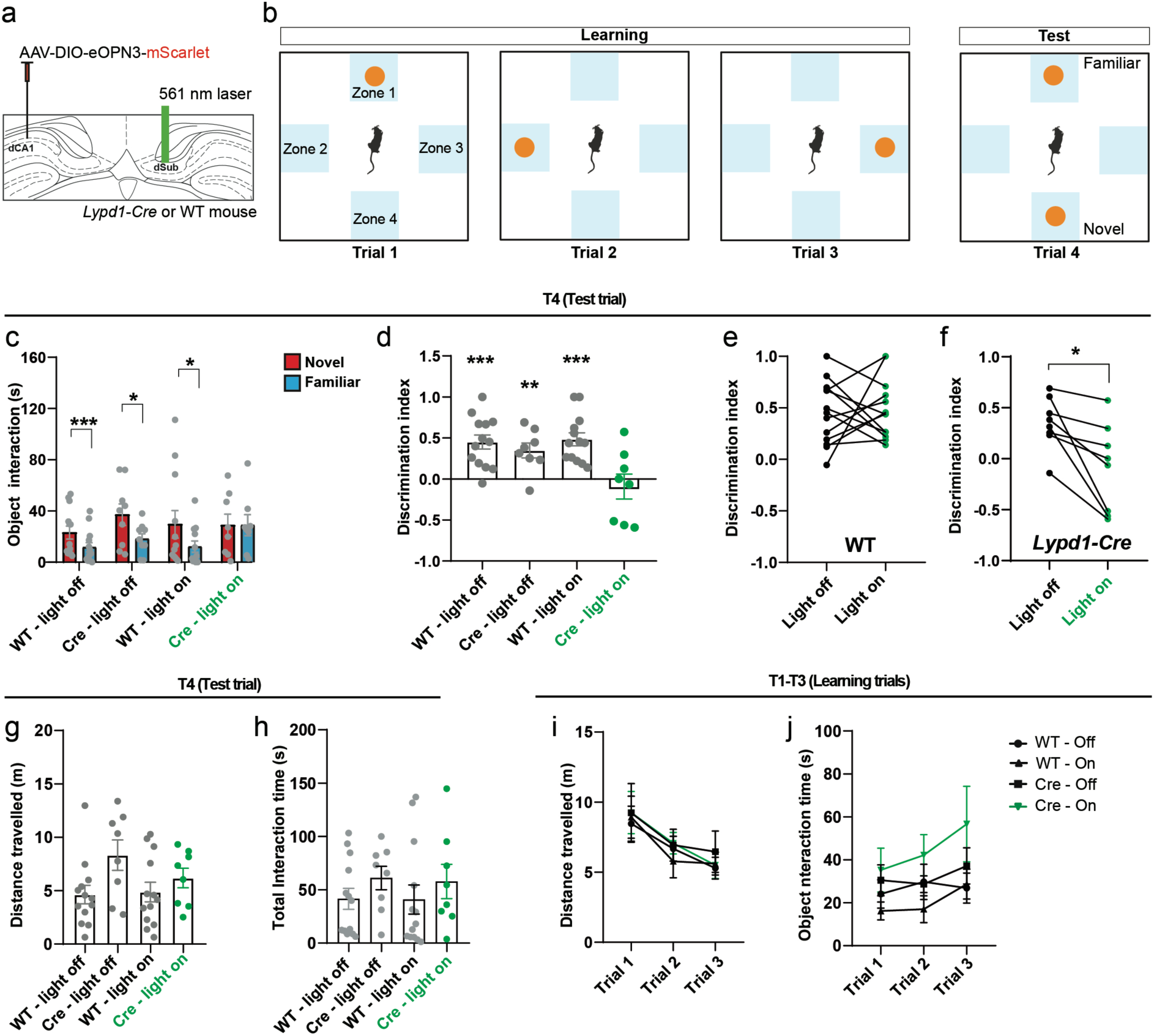
dCA1 projection to contralateral subiculum is necessary for spatial memory. **a.** *Lypd1-Cre* or WT mice from both sexes injected in the right dCA1 with AAV2/1 hSyn1-DIO-eOPN3-mScarlet-WPRE and implanted with an optic fiber above the left dSUB to silence dCA1 to dSUB interhemispheric terminals. **b.** Schematic of the object location test of spatial memory. **c.** Time of interaction with the object located in the familiar or novel position. **d.** Discrimination index for the novel vs. familiar location. **e.** Paired discrimination index for the novel location versus familiar in each WT mouse with and without light. **f.** Paired discrimination index for the novel versus familiar location in each *Lypd1-Cre* mouse with and without light. **g**. Total distance traveled during test trial. **h**. Total interaction time with objects during T4. **i.** Distance traveled during learning trials (T1-T3). **j**. Total interaction time with the object during learning trials (T1-T3). For the entire figure: bar graphs represent mean ± SEM and each point represents one mouse (13 WT and 8 *Lypd1-Cre* mice).

Because memory performance can be affected by anxiety, we examined the effect of silencing dCA1 projection to contralateral dSUB (Fig. S7a) in the open field (Fig. S7b) and elevated plus maze test (EPM, Fig. S7g). During the open field test, we found no effect on the total distance traveled, the time spent in center vs. surround (Fig. S7c-e) or in the number of entries into the center (Fig. S7f). Similarly, the EPM test yielded no differences in the distance traveled or the time spent in the open compared to the closed arms (Fig. S7h-j). In addition, we did not detect any difference in the number of entries to the open arm between WT and *Lypd1-Cre* mice. (Fig. S7k). Thus, silencing contralateral dCA1 projections to dSUB does not increase anxiety. Finally, as dorsal hippocampus and dorsal subiculum have been linked to object recognition,^20,69^ we also performed the novel object recognition test. We exposed the mice to two identical objects located in opposite locations within the open field during three consecutive trials to habituate them to the objects (learning phase: 1^st^ to 3^rd^ trial, Fig. S8a-b).^66^ During the test phase (4^th^ trial) we substituted one of the familiar objects with a new one. During the test phase all experimental groups spent more time with the novel object compared to the familiar one, indicating normal object novelty detection (Fig. S8c-d) The distance travelled and time of interaction with the objects during the learning or test phases were similar between all groups (Fig. S8e-h). Altogether these experiments suggest that dCA1 projection to contralateral dSUB regulates spatial memory without prominent effect on anxiety or novel object recognition.

### Interhemispheric projections from dCA1 to dorsal subiculum support spatial working memory

We then tested whether this projection was important for spatial working memory using the spontaneous alternation T-maze test (Fig. 3a-b).^65^ In this test, mice explore a T-shaped maze and have the option to enter the left or the right arm of the maze (Fig. 3b). Each mouse ran 6 consecutive trials and neither arm contained a reward. In control conditions, mice alternate between arms which is reflected as a percentage of alternation above chance (50%). The three control groups of our experiment alternated between arms above chance level (Fig. 3c), indicating normal working memory functioning. Silencing the projection (*Lypd1-Cre* mice with light on) brought the percentage of alternation to chance levels (Fig. 3c). We also compared the light on/off conditions within each mouse. Turning on the light decreased the percentage of alternation in *Lypd1-Cre* mice but not in WT (Fig. 3d-e). Finally, we measured the latency for the mouse to enter one arm or the other across the 6 consecutive trials. Contrary to the control mice that systematically chose quickly, the test mice needed much more time to decide which arm to visit starting from trial 2 (Fig. 3f), this increase in latency was also evident when comparing the mean time for all trials between groups (Fig. 3g). These experiments show that the interhemispheric projections from dCA1 to the dorsal subiculum regulate spatial working memory in addition to spatial memory.

**Figure 3:**
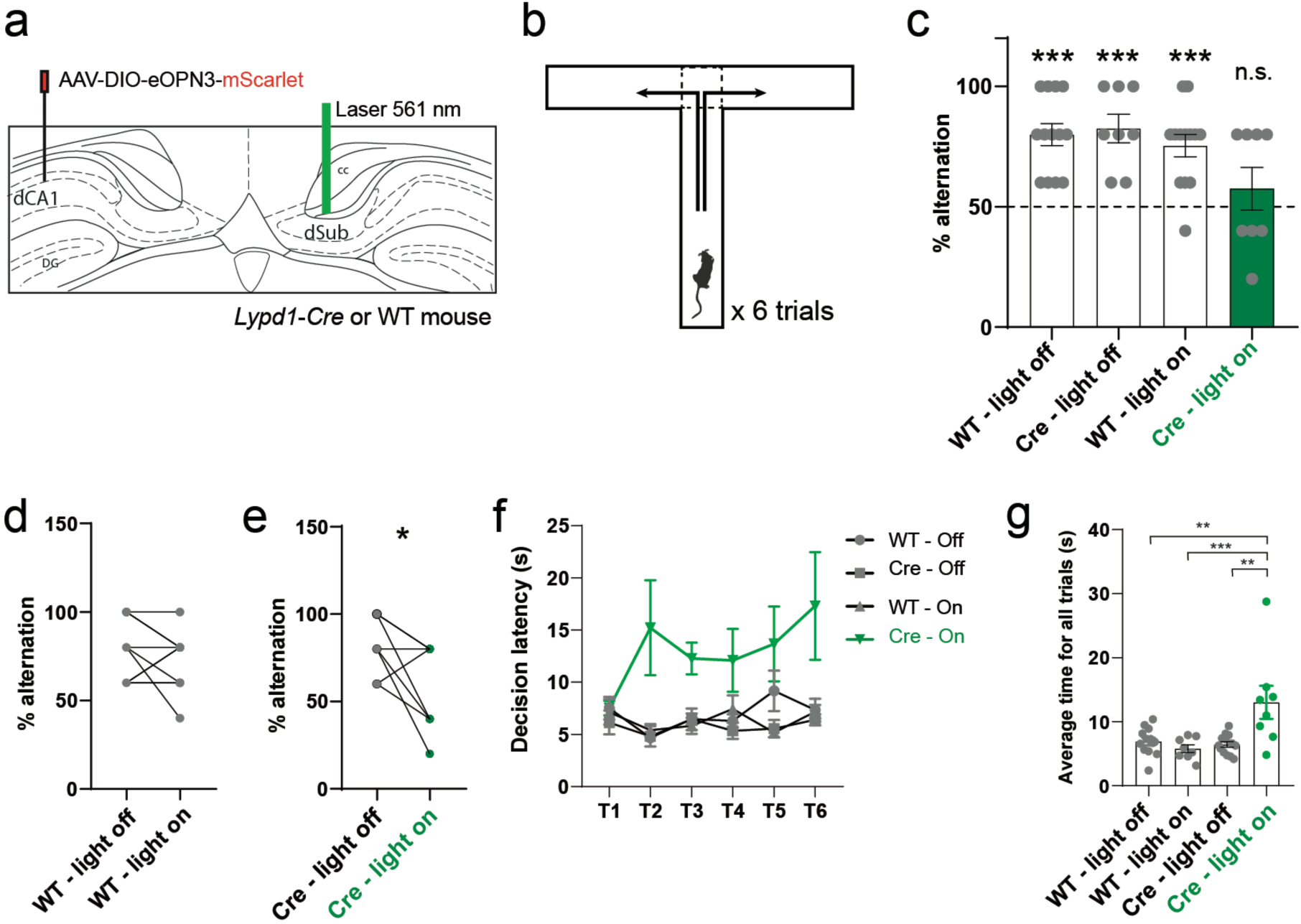
dCA1 projection to contralateral subiculum is necessary for spatial working memory. **a.** *Lypd1-Cre* or WT mice from both sexes injected in the right dCA1 with AAV2/1 hSyn1-DIO-eOPN3-mScarlet-WPRE and implanted with an optic fiber above the left dSUB to silence dCA1 to dSUB interhemispheric terminals. **b.** Schematic of the spontaneous alternation T-maze test for spatial working memory. **c.** Percentage of alternations in each group during the 6 consecutive trials. Each point corresponds to one mouse (13 WT and 8 *Lypd1-Cre* mice). **d.** Paired percentage of alternations in each WT mouse with or without light. **e.** Paired percentage of alternations in each *Lypd1-Cre* mouse with or without light. **f.** Decision latency (time spent before entering one arm) in each trial. **g**. Average decision time for all trials (T1-T6) in each group.

### Spatial cognition of male and female *Df16(A)^+/-^* mice is differentially impaired

As previous work reported altered cognition and dysregulated CA1 in *Df16(A)^+/-^* mice,^52–55,70^ we tested whether spatial memory and spatial working memory was impaired in male and female *Df16(A)^+/-^* mice using the same tests. To this end, we performed the object location test of spatial memory^64^, the spontaneous alternation T-maze test for spatial working memory^65^ and the novel object recognition test^66^ on female and male *Df16(A)^+/-^* mice and their WT littermates. In the object location test of spatial memory, we observed that only male mutants failed discriminate between a novel or familiar position (Fig. 4a-c), indicating disrupted spatial memory. Consequently, when comparing the discrimination indexes from WT and mutant mice, we found that the decrease was significant for males only. This change was not due to a change in locomotion as reflected by the distance traveled during the test (Fig. 4d), nor due to a lack of object interaction (Fig. 4e). Similarly, we did not observe any difference between mutant and WT during the learning phase (Fig. 4f-g). Then, mice performed the spontaneous alternation T-maze test (Fig. 4h).^65^ When analyzing the mice performance during this test, we observed that only mutant males failed to exhibit spontaneous alternation (Fig. 4i). However, despite mutant females showing a similar percentage of alternation compared to their WT littermates (Fig. 4i), both male and female mutant mice exhibited a higher decision latency before entering one arm of the maze or the other (Fig. 4j). This difference became even more evident when analyzing the average decision latency for all trials between groups (Fig. 4k). Finally, we also performed the novel object recognition test (Fig. S9a-b).^66^ Male and female *Df16(A)^+/-^* mice showed intact discrimination between a novel and a familiar object. (Fig. S9c-h). Overall, these results show that spatial cognition is impaired in *Df16(A)^+/-^* mice with male mutant mice being markedly more affected than females.

**Figure 4.**
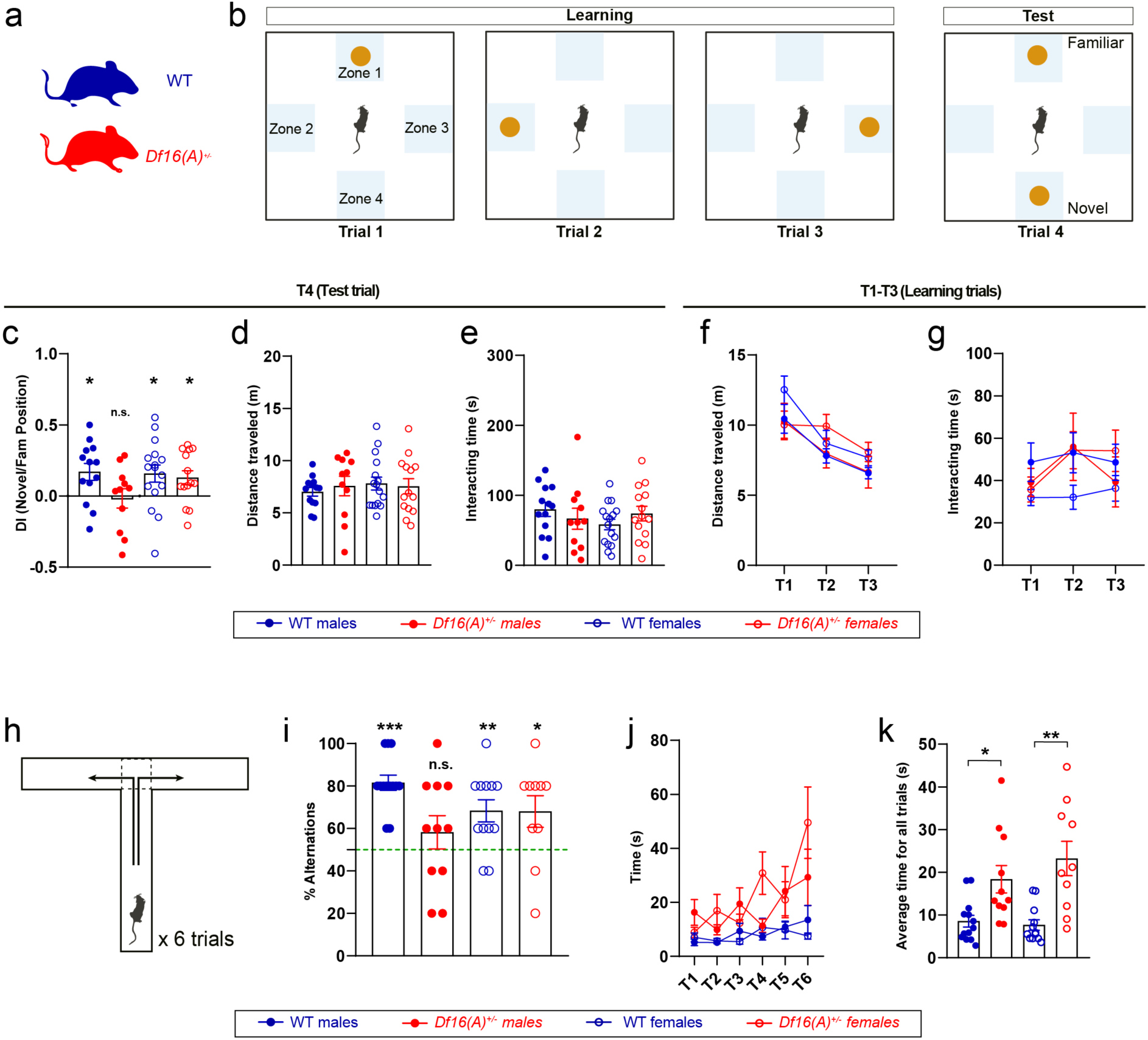
Spatial cognition of male and female *Df16(A)^+/-^* mice is differentially impaired. **a.** *Df16(A)^+/-^* or WT female and male mice were tested. **b.** Schematic of Object location test of spatial memory. **c.** Discrimination index for the novel vs. familiar location. In (c-e) each point represents one mouse (13 WT males, 11 *Df16(A)^+/-^* males, 16 WT females and 14 *Df16(A)^+/-^* females). **d.** Total distance traveled during test trial. **e**. Total interaction time with objects during the entire test. **f.** Distance traveled during learning trials (T1-T3). **g**. Total interaction time with the object during learning trials (T1-T3). Schematic of the T-maze test of spontaneous alternation for spatial working memory. **i.** Percentage of alternations in each group during the 6 consecutive trials. In (i-k), each point corresponds to one mouse (13 WT males, 11 *Df16(A)^+/-^* males, 12 WT females, and 10 *Df16(A)^+/-^* females). **j.** Decision latency (time spent before entering one arm) in each trial. **k.** Average decision latency for all trials. For the entire figure, bar graphs represent mean ± SEM.

### Interhemispheric projections from dCA1 to the contralateral hippocampal formation are differentially disrupted in male and female *Df16(A)^+/-^* mice

Since *Df16(A)^+/-^* mice exhibit dysregulation of their dCA1^52–54^ and deficits in CA1-dependent behaviors such as spatial working memory^55^ or fear conditioning,^48^ we investigated whether a decrease in interhemispheric inputs from CA1 to subiculum could contribute to the spatial cognition defects exhibited by *Df16(A)^+/-^* mice.^55,70^ In addition, we asked whether the sex-specific changes in spatial cognition we observed could be corelated with anatomical differences between male and female mutants. We injected the retrograde agent CtB-488 in dCA1 of the right hippocampus of adult female and male *Df16(A)^+/-^* mice and control littermates before imaging (Fig. 5a-b). We only kept brains with similar CtB-488 injection sites (Fig. S10a) and counted the number of retrogradely labelled cells in contralateral (left) dCA1, dCA2 and dCA3. We did not find any difference in the number of CtB^+^ cells between WT and mutant male mice in dCA2 and dCA3 (Fig. S11a-d), suggesting that the interhemispheric projections originating from these regions are not altered in *Df16(A)^+/-^* male mice. In dCA1 however, *Df16(A)^+/-^* male mice exhibited a marked decrease in CtB^+^ cells in distal, intermediate and proximal dCA1 (Fig. 5c-d). Injections in female mice revealed a decrease in CtB^+^ cells only in contralateral distal dCA1 but not in intermediate or proximal regions (Fig. 5e-f & Fig. S11e-h). As previous publications have shown the remarkable differences between deep and superficial neurons of the pyramidal CA1 layer,^71–74^ we further analyzed whether the reduction originated preferentially from deep or superficial layers and found that both layers contribute equally to the decrease (Fig. S12a-c). We conclude that *Df16(A)^+/-^* mice exhibit a decrease in dCA1 projection to contralateral dCA1 which is more pronounced in male mice where all CA1 regions are affected compared to females where defects are limited to distal dCA1 only.

**Figure 5.**
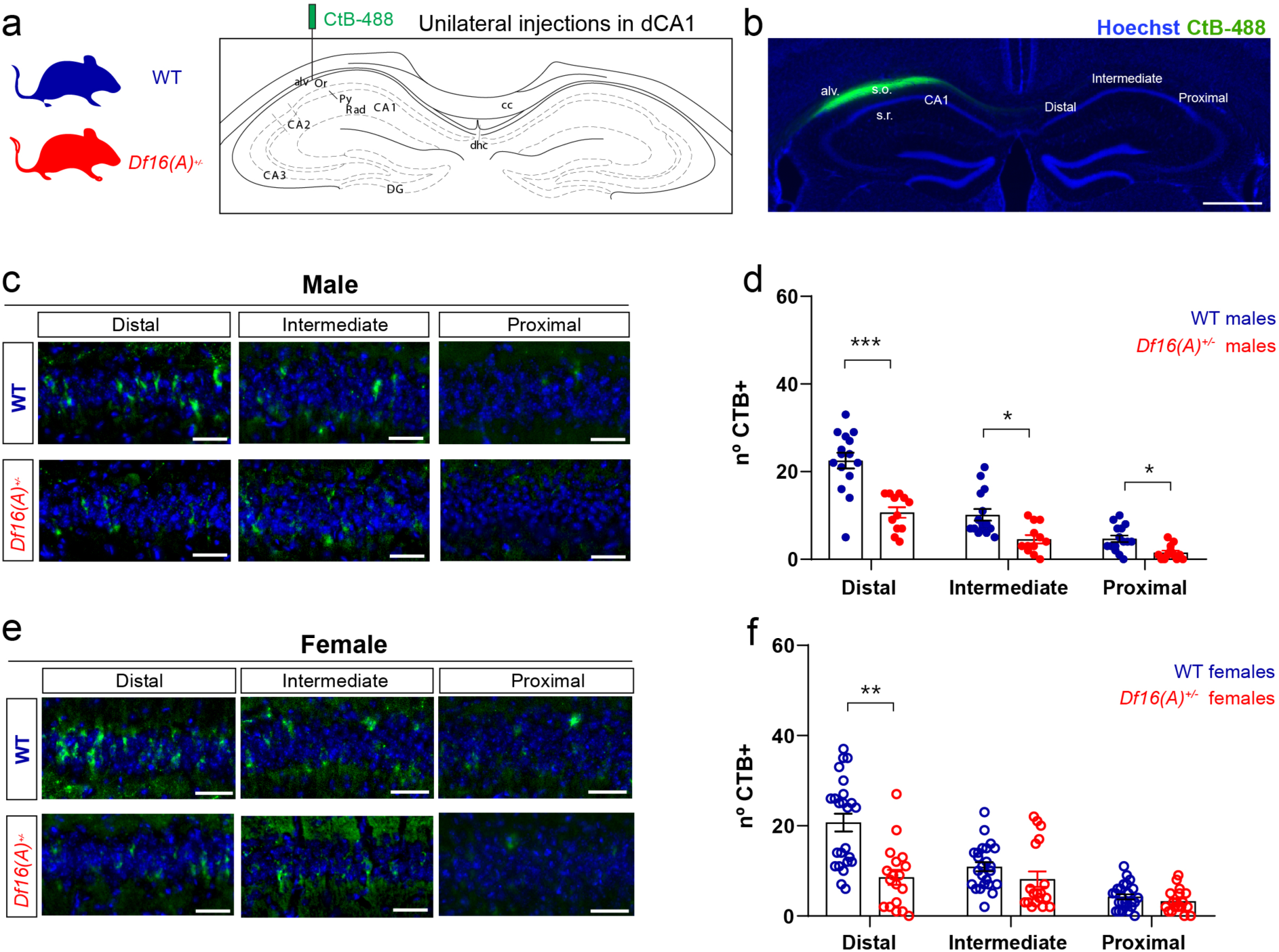
dCA1 projections to contralateral dCA1 are dysregulated in *Df16(A)^+/-^* mice. **a.** Injection of CtB-488 in the right dCA1 of *Df16(A)^+/-^* mice and littermates. **b.** Coronal section showing injection site and diffusion of CtB-488 in the right dCA1. **c.** Representative images of CtB^+^ cells in distal, intermediate and proximal contralateral dCA1 (left CA1) from *Df16(A)^+/-^* male mice and littermates. **d.** Number of CtB^+^ cells in distal, intermediate and proximal contralateral dCA1. Each point corresponds to one observation (5 WT and 4 *Df16(A)^+/-^* mice, 3 observations per mouse). **e.** Representative images of CtB^+^ cells in distal, intermediate and proximal contralateral dCA1 from WT and *Df16(A)^+/-^* female mice. **f.** Number of CtB^+^ cells in distal, intermediate and proximal contralateral dCA1 from WT and *Df16(A)^+/-^* male mice. Each point corresponds to one observation (8 WT and 6 *Df16(A)^+/-^* mice, 3 observations per mouse). For the entire figure, bar graphs represent mean ± SEM. Scale bars: 500 µm (b) and 50 µm (c,e).

To evaluate whether dCA1 projections to contralateral dSUB are also be dysregulated in *Df16(A)^+/-^* mice, we injected CtB-488 into the dSUB of the right hemisphere of male and female *Df16(A)^+/-^* mice and control littermates and counted CtB^+^ cells in the left dCA1(Fig. 6a-b). As previously, we only analyzed brains with comparable CtB-488 injection sites in dSUB (Fig. S10b). Mutant male mice exhibited a decrease in CtB^+^ cells specifically located in the contralateral distal but not medial or proximal dCA1 (Fig. 6c-d). This decrease was equally present in deep and superficial layers (Fig. S12 d-f). In females *Df16(A)^+/-^* mice, we found no significant difference in the number of CtB^+^ cells between WT and *Df16(A)^+/-^* females despite a tendency for a decrease in distal CA1 CtB^+^ cells (Fig. 6e-f). Overall, these experiments show that dCA1 interhemispheric projections to the contralateral formation are preferentially impaired in male compared to female mutant and suggest that disruption of the interhemispheric connectivity may lead to the deficit in spatial cognition exhibited by *Df16(A)^+/-^* mice.

**Figure 6:**
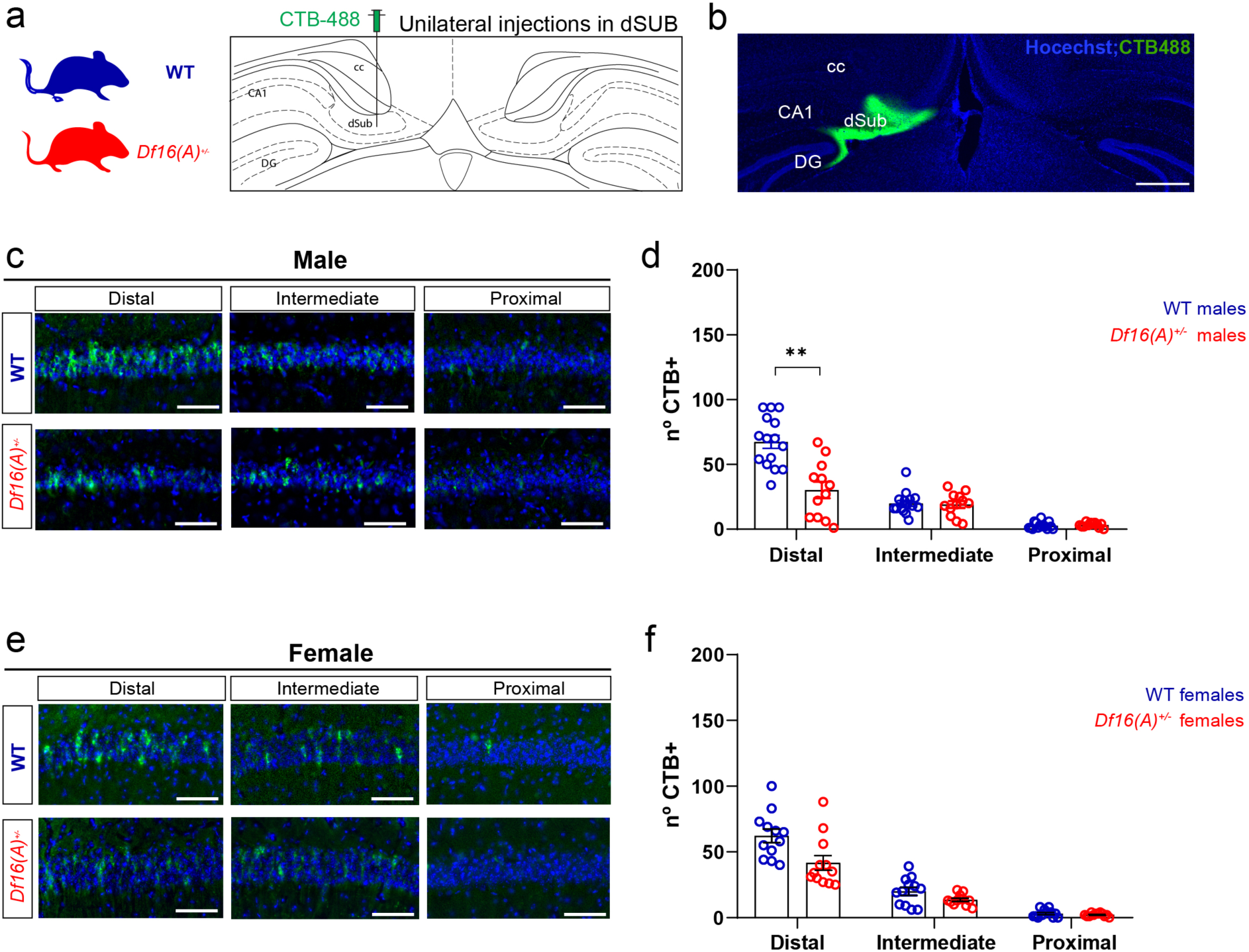
dCA1 interhemispheric projections into contralateral dSUB are dysregulated in *Df16(A)^+/-^* mice. **a.** WT and *Df16(A)^+/-^* mice injected with CtB-488 in the right dSUB. **b.** Coronal section showing injection site and diffusion of CtB-488. **c.** Representative images of CtB^+^ cells in distal, intermediate and proximal contralateral dCA1 of WT and *Df16(A)^+/-^* male mice. **d.** Number of CtB^+^ cells. Each point represents one observation (5 WT and 4*Df16(A)^+/-^* mice, 3 observations per mouse). **e.** Representative images of CtB^+^ cells in distal, intermediate and proximal contralateral dCA1 of WT and *Df16(A)^+/-^* female mice. **f.** Number of CtB^+^ cells. Each point represents one observation (4 WT and 4 *Df16(A)^+/-^* mice, 3 observations per mouse). For the entire figure, bar graphs represent mean ± SEM. Scale bars: 500 µm (b) and 50 µm (c,e).

## Discussion

The ipsilateral connectivity of the hippocampus has been extensively studied to understand the neurobiology of complex cognitive processes such as learning and memory or spatial navigation. However, we still know very little about the detailed organization of interhemispheric hippocampal connections which is crucial as hippocampal information processing relies on a dynamic exchange of information between both hemispheres. Here, we show how dCA1 pyramidal neurons send interhemispheric projections to several brain regions within and outside the hippocampal formation. We found interhemispheric projections to dorsal CA1 and dorsal subiculum. Within the hippocampus proper, dCA1 interhemispheric projections target preferentially the stratum radiatum of contralateral CA1 unlike interhemispheric projections from dCA3 and dCA2 which innervate the stratum radiatum and stratum oriens of contralateral CA1 to the same extent.^75^ Whether projections from dCA1 or dCA3 target different neurons in contralateral dCA1 remains unknown but the different projection patterns suggests a functional segregation. CA1 pyramidal neurons also innervate the contralateral subiculum in a very defined pattern, with axons only visible in the dorsal subiculum region (also known as subiculum proper) but not in the adjacent pro-subiculum or ventral subiculum regions. Our retrograde tracing experiment showed that the majority of dCA1 neurons projecting to the contralateral dorsal subiculum had their somas located in the distal part of CA1, a region which plays a prominent role in spatial navigation due to its preferential MEC inputs.^76,77^

To a lesser extent, we also visualized contralateral projections from dCA1 to the intermediate lateral septum (LSI), laterodorsal nucleus of the thalamus (LDDM), reuniens nucleus (Re) and rhomboid nucleus (Rh). LSI neurons harbor spatial information^78^ and the LDDM has been shown to play a critical role in spatial learning and memory.^79^ How these interhemispheric collaterals participate in shaping the spatial information conveyed to the LSI and LDDM remains unknown. Given the extensive lateralization of the hippocampus,^11^ it is likely important that ipsi- and contralateral information are integrated in downstream targets. The rhomboid and reuniens nuclei form reciprocal connections with dCA1,^80,81^ which play a role in perception and cognition^82–84^ but, here as well, the contribution of the interhemispheric collaterals to this network are unknown. We also reported that CA1 axons cross the midline through both the dorsal and ventral commissures. Traditionally, the ventral hippocampal commissure has been linked to bilateral integration of information carried by neurons within the hippocampus proper, while the dorsal commissure also exhibit interhemispheric fibers from the para-hippocampal formation.^20^ Therefore, we can speculate that CA1 pyramidal neurons cross the midline through one or the other commissure depending on the spatial location of their interhemispheric targets and function.

The function of the interhemispheric projection from dCA1 to contralateral dCA1 was previously characterized by Zhou et al.,^19^ showing its importance in the generalization process occurring following fear memory acquisition. We therefore focused on the function of the interhemispheric projection from dCA1 to dorsal subiculum and proved this projection to be necessary for spatial memory and spatial working memory. These results expand on previous studies demonstrating the importance of subiculum cells for spatial memory.^85–87^ Importantly, silencing the projection had no effect on exploration, locomotion or anxiety which aligns with the proposed role for the dorsal subiculum in processing spatial information while the ventral subiculum regulates anxiety, mood and emotions.^69^

In support of our observation that interhemispheric projection of dCA1 pyramidal neurons supports key hippocampal-dependent cognitive functions in WT mice, our characterization of these projections in a mouse model of 22q11.2 deletion, revealed that dCA1 contralateral projections are disrupted by a mutation predisposing to cognitive dysfunction and SCZ. We observed that, while male *Df16(A)^+/-^* mice exhibit a decrease in the CA1 interhemispheric projection targeting contralateral dCA1 and dorsal subiculum, female *Df16(A)^+/-^* mice only showed minor alterations in the dCA1-to-dCA1 circuit. Moreover, the dCA1-to-dCA1 reduction was less severe in females than in males. Characterization of female and male *Df16(A)^+/-^* mice cognition revealed impairments in spatial memory and spatial working memory only in male mutants. This may be due to the more pronounced disruption of dCA1 contralateral projections observed in male *Df16(A)^+/-^* mice. Our results are consistent with a recent functional magnetic resonance imaging study of a similar 22q11.2DS mouse model (LgDel mice), which revealed age-specific patterns of functional dysconnectivity within the hippocampal formation, with widespread hyper-connectivity in juvenile mice reverting to focal hippocampal hypoconnectivity over puberty.^47^ Analysis of both 22q11.2DS mouse models therefore points to a decrease in hippocampal connectivity in the adult brain. Our results are also consistent with a number of studies in 22q1.2DS patients and mouse models that indicate sexual dimorphism in cognitive impairment with males more affected than females.^48,88^

Despite evidence for early alteration in hippocampal connectivity in 22q11.2DS patients and mouse models, the cause(s) for these changes remains unknown. It is possible that hemizygosity of one or more genes within the microdeletion (such as *ZDHHC8* ^89^) induces defects in axonal growth in hippocampal neurons from *Df16(A)^+/-^* mice during embryogenesis which leads to altered arborization and synapse maturation. An alternative, intriguing, possibility relates to the observation that CA1 neurons from *Df16(A)^+/-^* mice present alterations in their excitatory/inhibitory input (E/I) balance,^52^ a ratio that is critical for stabilization of interhemispheric projections during development. For example, a change in the E/I ratio of developing L2/3 neurons of the cortex, which are a major interhemispheric population, is sufficient to disrupt their connectivity to the contralateral hemisphere.^90^ In developing CA1 pyramidal neurons, the E/I ratio is likely to change due to changes in their dendritic tree and spines as well as electrophysiological properties.^47,48,52^ Indeed, a recent study of CA1 interneurons in adult *Df16(A)^+/-^* mice found that the while the density of various interneuron types is unchanged, their activity is markedly disrupted during random foraging and goal-oriented reward learning tasks.^53^ Our results combined with these other studies therefore raises the possibility that an imbalance in E/I inputs to CA1 neurons occurring during the development of 22q11DS mice and patients may structurally disrupt CA1 interhemispheric projections and impair cognition in the adult. Whether changes in projection are due to activity dependent mechanisms^47^ or not is still unclear and future investigations, taking into account the sexual dimorphism of the microdeletion phenotype, are required to decipher the underlying cellular mechanisms.

### Technical limitations of our study

Even though the previous characterization of the *Lypd1-Cre* line showed equivalent CRE expression between deep and sup CA1 layers,^57^ in our hands, most of the cells expressing the GFP were in the deep layer. We presume this might be due to the fact that the virus was injected on the dorsal side of the pyramidal layer. It is also possible that the AAV serotypes we used have a tropism for the deep pyramidal neurons. In our retrograde tracing experiments performed from the dCA1 and dSUB we did not detect a difference in the number of contralateral CTB^+^ cells between deep and sup layers, suggesting that both populations equally contribute to the contralateral targeting of dCA1 and dSUB. However, we cannot discard the possibility that some superficial CA1 neurons project to additional regions in the contralateral hemisphere. Then, the limited CtB uptake prevents us from quantifying the absolute number of CA1 neurons projecting to the contralateral subiculum of dCA1. It also prevents us from investigating whether the same CA1 pyramidal neuron bifurcates toward several targets or whether there is different population of dCA1 pyramidal neurons. However, dCA1 neuron axons are known to have few or no collaterals.^91^ Finally, we traced and probed the function of the right CA1 to the left subiculum. Future studies should consider a possible lateralization of this interhemispheric projection and probe the projection and function from the left dCA1 to the right subiculum.

## Supporting information

Statistical tables

## Acknowledgements

We thank Antoine Besnard, Jorge Brotons Mas, Raul Andero and Larry Swanson for comments on the manuscript. This project has received funding from the European Research Council (ERC) under the European Union’s Horizon 2020 research and innovation programme (grant agreement No 949652 to F.L.). F.L. also acknowledges support from CIDEGENT grant from the Valencian Community and the Severo Ochoa Foundation. N.S.L.R was funded by a Severo Ochoa fellowship from the Spanish MINECO (BES-2015-071690). M.N. acknowledges PID2020-112831GB-I00 funded by MCIN/AEI /10.13039/501100011033. J.A.G. acknowledges support from NIMHR01MH124047 grant. The project that gave rise to these results received the support of a fellowship from “la Caixa” Foundation (ID 100010434) to M.H.B-G. The fellowship code is LCF/BQ/DI20/11780018.

## Author contributions

Conceptualization: N.S.L.R., M.N. J.A.G and F.L.

Investigation: N.S.L.R., M.H.B-G. and M.J.G

Viral vector cloning: C.G.F.

Writing – original draft: N.S.L.R. and F.L.

Writing – review & editing: N.S.L.R, M.N., J.A.G. and F.L.

Visualization: N.S.L.R.

Supervision: M.N. and F.L.

Funding acquisition: M.N. and F.L.

## Materials and Methods

### Key resources

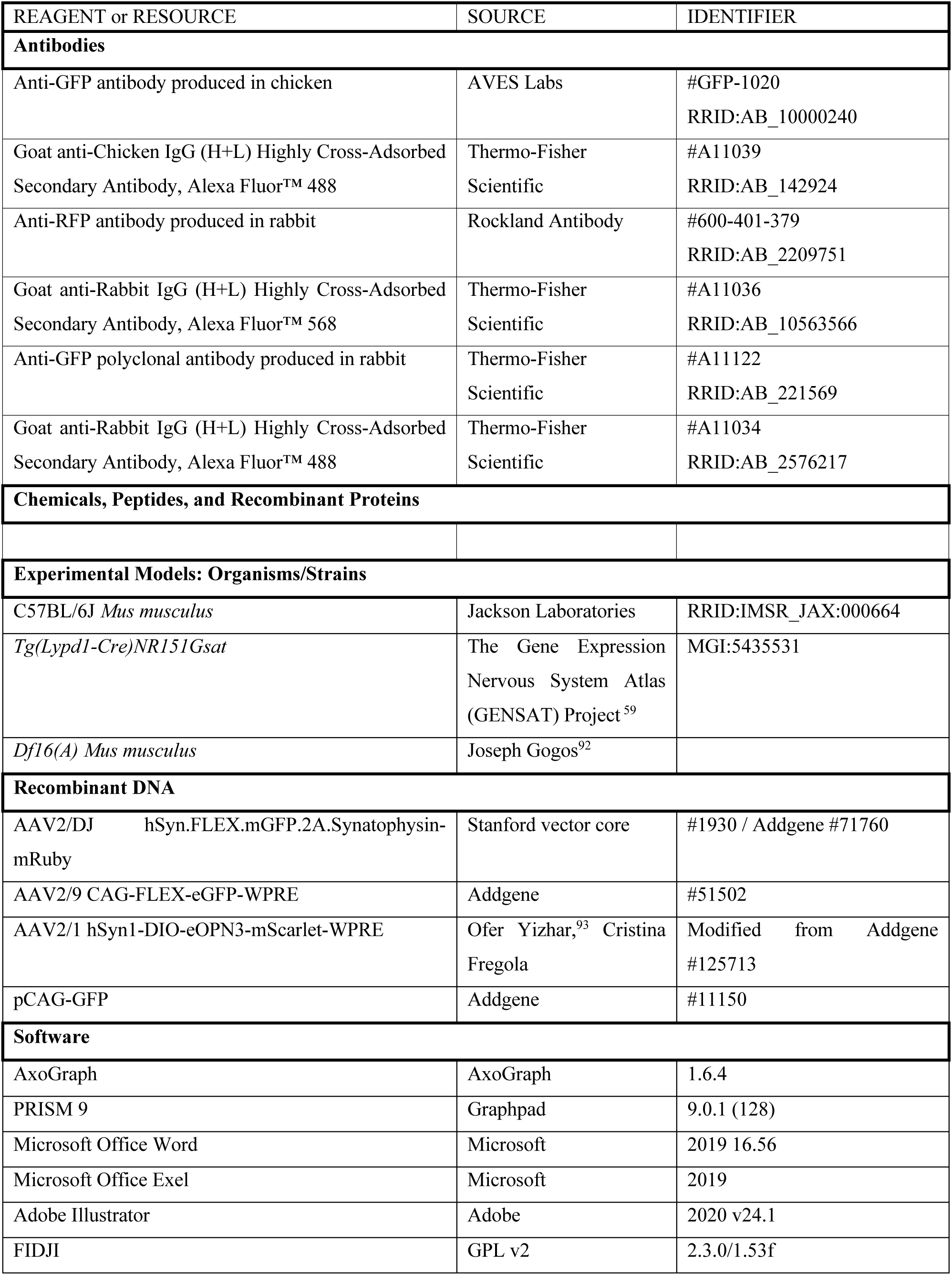
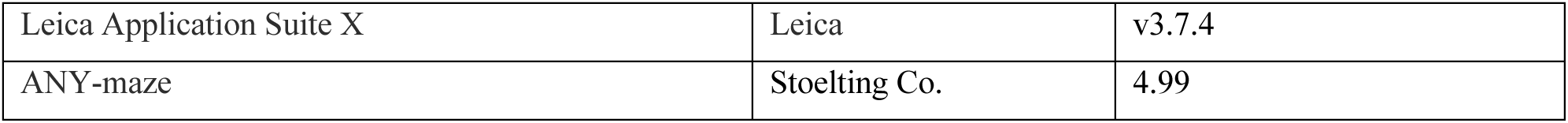

### Experimental subjects

Animal procedures were approved by the CSIC, the Community of Madrid Ethics Committees on Animal Experimentation and the Generalitat Valenciana Agriculture Department in compliance with national and European legislation. We used P7 to 16-week-old C57BL6/J wild-type (Jackson Laboratories, #000664) mice as well as 2-to 4-month-old mice from the following transgenic mouse lines: *Lypd1-Cre* mice (Tg-Lypd1-Cre, NR151Gsat MGI:5435531)^57^ and *Df16(A)* mice^92^. All transgenic mice were maintained on the C57BL6/J background. For experiments developed at early developmental stages, the day we observed a vaginal plug was defined as embryonic day E0.5. Animals were housed and maintained following the guidelines from the European Union Council Directive (86/609/ European Economic Community). All the procedures for handling and sacrificing animals followed the European Commission guidelines (2010/63/EU). All animal procedures were approved by the CSIC and the Community of Madrid Ethics Committees on Animal Experimentation in compliance with national and European legislation (PROEX).

### Generation of the conditional eOPN3/Scarlett-expressing AAV vectors

To subclone the coding sequence of eOPN3-Scarlet into a double-floxed inverted open-reading frame AAV vector, an EcoRI-BamHI fragment, containing the eOPN3-Scarlet open reading frame (ORF), was removed from pAAV-hSyn-SIO-eOPN3-mScarlet-WPRE (Addgene 125713), blunted and ligated into the NheI and AscI digested, blunted and dephosphorylated pAAV-hSyn-DIO-mCherry (Addgene 50459) vector. The plasmid was then packaged into AAV9 viral vectors at the Neurotropic Vector Unit of the Instituto de Neurociencias following Addgene protocols.

### Virus injections

For all injections, animals were anesthetized using isoflurane and given analgesics before surgery. A craniotomy was performed above the region of interest and a glass pipette was stereotaxically lowered down to the desired depth. All injections were performed with a nano-inject II (Drummond Scientific). Pulses of 9.2 nl were delivered 10 seconds apart until the total amount was reached. 2 minutes after infusion of the entire volume, the pipette was slowly retracted. For CA1 interhemispheric projections tracing we injected 200 nl of AAV2/DJ hSyn.FLEX.mGFP.2A.Synatophysin-mRuby (Addgene, #71760, prepared by the Stanford University vector core #1930). For specific silencing of CA1 interhemispheric terminals in dSUB we injected 200 nl of AAV2/1 hSyn1-DIO-eOPN3-mScarlet-WPRE (Addgene 125713-AAV1). For retrograde tracing from dCA1 or dSUB we injected 150 nl of CtB-488 at 0.5% (Life Technologies, #C22841). dCA1 injection coordinates were the following from Bregma: antero-posterior -2.2 mm, medio-lateral +1.3 mm, dorso-ventral +1.7 mm. dSUB injection coordinates: antero-posterior -2.69 mm, medio-lateral +0.9 mm, dorso-ventral -2.15. vCA1 injection coordinates: antero-posterior -3.07 mm, medio-lateral +3.2 mm, dorso-ventral -2.75 mm. One week after CtB injection or after 2-3 weeks of virus expression, mice were anesthetized using isoflurane, perfused with 0.9% saline and their brains were quickly extracted and incubated in 4% paraformaldehyde (PFA) overnight for posterior immunohistochemical analysis.

### In utero electroporation

In utero electroporation was performed as previously described to restrict the neuronal population carrying a reporter gene (GFP). The transfection of the neuronal population of interest is achieved by developing DNA microinjections into the lumen of the ventricular system of mice embryos. We temporally restrict the expression of our reporter by developing the electroporation at E14, the moment in which most pyramidal neurons of the hippocampus are born. Spatially, we restrict the expression of the reporter based on the generation of an electric field, that will favor the transfection of the negatively charged DNA into the selected cell population.^94^ Precursors of CA1 pyramidal neurons were targeted by placing the two positive electrodes at both sides of the brain while placing the negative electrode above the brain forming an angle of 90 or 30° respective to the two positive electrodes in order to target the hippocampal precursors of all hippocampus or CA1 respectively. For the surgery, timed pregnant mice were anesthetized with isoflurane/oxygen. After exposing the embryos, we injected a solution of 1μg/μl containing the plasmid pCAG-GFP (Addgene plasmids #11150) into the embryo’s lateral ventricle using a 30 μm pulled glass micropipette. Five voltage pulses (36 mV, 50 ms) were applied using three external paddles oriented to target CA1 specifically or the whole hippocampus.^61^ After birth, brains were fixed by intracardiac perfusion at P7 for posterior immunohistochemical analysis.

### Optical ferrule implants

Animals were anesthetized using isoflurane and given analgesics. The skin above the skull was removed and a craniotomy was performed above the target region. Then the optical ferrule was lowered until the desired depth. Superglue was applied to hold the lens in position and then dental cement (GC FujiCEM 2) was applied to cover the exposed skull and keep the optical ferrule in position. Animals were allowed to recover for 5 days before being used. For silencing dCA1 projection to contralateral dSUB, we implanted optical ferrules with a core diameter of 200 µm of diameter, 2 mm of length and 0.39 numerical aperture (Neurophotometrics, R-Foc-L200c-39NA) in the left dSUB at the following coordinates from Bregma: antero-posterior -2.6 mm, medio-lateral +0.87 mm, dorso-ventral +1.7 mm.

### Immunohistochemistry

#### Adult brains

For labelling mCherry and GFP in adult brains, mice were anesthetized using isoflurane, perfused in the heart with 0.9% saline and their brains quickly extracted and incubated in 4% PFA overnight at 4°C. The day after, brains were washed for 1 hour in a solution of PBS with 0.3 M glycine. Adult brains were cut using a vibratome VT1000S (Leica Biosystems). Unless indicated otherwise, slices were permeabilized for 1h in PBS with 0.5% Triton-X100 (T9284, Sigma-Aldrich) before being incubated overnight at 4°C with primary antibodies diluted in PBS containing 0.5% Triton-X. Then, the slices were washed in PBS for 1 hour and then incubated overnight at 4°C with secondary antibodies from Thermo-Fisher Scientific at a concentration of 1:500 diluted in PBS with 0.1% Triton-X. For labelling mCherry and GFP in adult brains, we used rabbit anti-RFP (1:500, Rockland Antibody, #600-401-379) and chicken anti-GFP (1:1000, Aves, #GFP-1020) primary antibodies. Secondary incubation was performed with anti-rabbit antibody conjugated to Alexa-568 (#A11036) and anti-chicken antibody conjugated to Alexa-488 (#A11039). Hoechst counterstain was applied (Hoechst 33342 at 1:1000 for 30 min in PBS at room temperature) prior to mounting the slice using fluoromount (Sigma-Aldrich).

#### P7 brains

For labelling GFP in P7 brains, mice were anesthetized using isoflurane, perfused in the heart with 4% PFA and then post-fixed in 4% PFA overnight. The next day, brains were changed into a solution of 30% Sucrose in PBS to favor cryopreservation. After cryopreservation, we made 50 µm cryosections in the cryostat (-16°C) for immunohistochemistry. Slices were incubated with the primary antibody in PBST 0.05% overnight at 4°C. The next day we developed 3×10 min PBS1X washes before secondary incubation. We did secondary incubation at room temperature for 45 min. We used the primary antibody rabbit anti-GFP 1:500 (#A11122) followed by anti-rabbit antibody conjugated to Alexa-488 (1:500, ThermoFisher Scientific, #A11034). Cell nuclei were counterstained with 4′,6-diamidino-2-phenylindole (DAPI, 1:1000, Sigma, #D9542) during 10 minutes in PBS1X. Images were acquired using inverted confocal microscopes (LSM 900, Zeiss and SPII, Leica) or an epifluorescent microscope (Thunder, Leica).

### Behavioral tests

Based on our experience conducting behavior experiments, we used 6-10 animals per group. Animals with viral expression outside the region of interest, or implants that were not properly placed in the region of interest were excluded from analysis. The observer was blind to the identity of the mice while performing the behavioral experiments and the subsequent analyses. For all tests, we automatically tracked the mice using the software Any-Maze 7 from Stoelting. For all tests, mice were randomly exposed to light on or light off condition first and then one week later to the other condition. We used the first cohort of mice for the open field and elevated plus maze tests (except for three mice that lost their implant before testing for the EPM). Then, we used a second cohort for the spatial novelty, object novelty, and spatial working memory tests. The first cohort was composed of n = 4 females and n = 6 males. The second cohort was composed of n = 7 females and n = 9 males. All tests were performed with at least 1 week of interval.

#### Open field test

Mice were introduced into a previously unknown arena (60 cm x 60 cm) and allowed to freely explore for 5 min. Using automatic tracking of the test mouse (Any-Maze 7, Stoelting), we quantified the total distance traveled as well as time spent in the surround (20% of the surface) or center (remaining 80% of the surface) of the arena.

#### Elevated plus-maze (EPM) test of anxiety

This test was performed using the EPM form Harvard apparatus designed for mice (#LE842A). Mice were placed at the center of a maze consisting of a cross with two open arms and two closed arms. They were allowed to explore the maze freely for 5 min. We quantified the amount of total distance traveled and the time and ratio spent in open or closed arms using automatic tracking of the test mouse (Any-Maze 7, Stoelting).

#### Object location test of spatial memory

This test was performed in the same arena as the open field test. During the learning phase, the mice were allowed to explore the arena with the object in it for 5 minutes. The learning phase consisted of 3 trials separated by 3 min intervals. In each trial, the object was placed at a different position within the arena. 30 minutes later the mice were placed again in the arena with the object back to its initial position and another identical object placed in a novel position. This last trial also lasted 5 minutes. We measured the time investigating each object during the last trial as well as the total distance traveled and object interaction time during 3 learning trials. For all tested mice, we followed the same order of spatial alternation as reflected in Figure 4.

#### Novel object recognition test

This test was performed in the same arena as the open field test. During the learning phase, the mice were allowed to explore the open field for 5 min with two identical objects placed at opposite corners. We repeated this trial three times with 3 minutes intervals. 30 minutes later the mice were introduced again into the open field but this time, one of the familiar objects was substituted with a novel. This test trial also lasted for 5 minutes. Throughout the experiment, we alternated the position in which we placed the novel object to avoid possible bias. We measured the time spent investigating the novel or familiar objects during the test trial, and total distance traveled and object interaction during the three consecutive learning trials.

#### Spontaneous alternation T-maze test for spatial working memory

We built a T-maze with white opaque polymethylmethacrylate following previously published measurements for the test: The T-maze was mounted 65 cm above the floor. The schematic of the apparatus is shown in Figure 4. All arms (the starting arm and the two goal arms) were designed 7 cm broad and 35 cm long. Therefore, the central choice was a region of 7 x 7 cm^2^. The test consisted of 6 consecutive trials with no interval, in which the mice were placed at the beginning of the starting arm (far from the central choice area). The mice traveled until the central choice where they decided to explore either the left or the right arm (goal arms). Once the mice entered the selected arm, we blocked the exit of the mice and allowed them to stay there for 30 seconds. After this time, the mice were placed again in the starting arm. We repeated this operation for 6 consecutive trials and annotated each time the explored arm and the latency to reach an end of the maze. We calculated the alternation score by dividing the number of alternations with the total number of trials. We follow all the indications of the protocol previously published^65^.

#### Optogenetic terminal silencing

Mice were habituated to the patch cord before testing their behavior. In the experimental condition of the silencing, light stimulation was developed using a 561 nm laser (LaserGlow) adjusted at 5 mW and applied during all the trials of the tests (learning and test phases). In control conditions, mice were subjected to the test with the patch cord connected but without light stimulation.

### Quantifications and statistical analysis

Statistical tests were performed using PRISM 9 (Graphpad) and the details of the test can be found in the figure legends. Results presented in the text, figures and figure legends are reported as the mean ± SEM. In all figures, * is for *p* < 0.05, ** is for *p* < 0.01, *** is for *p* < 0.001. When multiple observations were done in the same mouse, we used nested statistical tests to consider the lower degree of freedom. We classified CtB^+^ cells as deep or superficial drawing a line halfway through the stratum pyramidale^74^. We used the Paxinos atlas (4^th^ edition) to delineate separations between brain regions^95^CA1. Each point corresponds to one observation (4 mice, 3 sections per mouse). Nested *t* tests, *p* < 0.0001. **f-n.** Coronal hippocampal sections showing GFP^+^ cells in anterior (f), medial (i) and posterior ipsilateral dCA1 (l) and GFP^+^ fibers and mRuby^+^ pre-synaptic terminals in contralateral dCA1 and dSUB. For the entire figure, bar graphs represent mean ± SEM. Scale bars: 1mm (b), 300 µm (f,g,i,j,l,m) and 100 µm (c,h,k,n). (alv.): alveus, (s.o): stratum oriens and (s.r.): stratum radiatum.

---

**Supplementary figure 1 related to figure 1.**
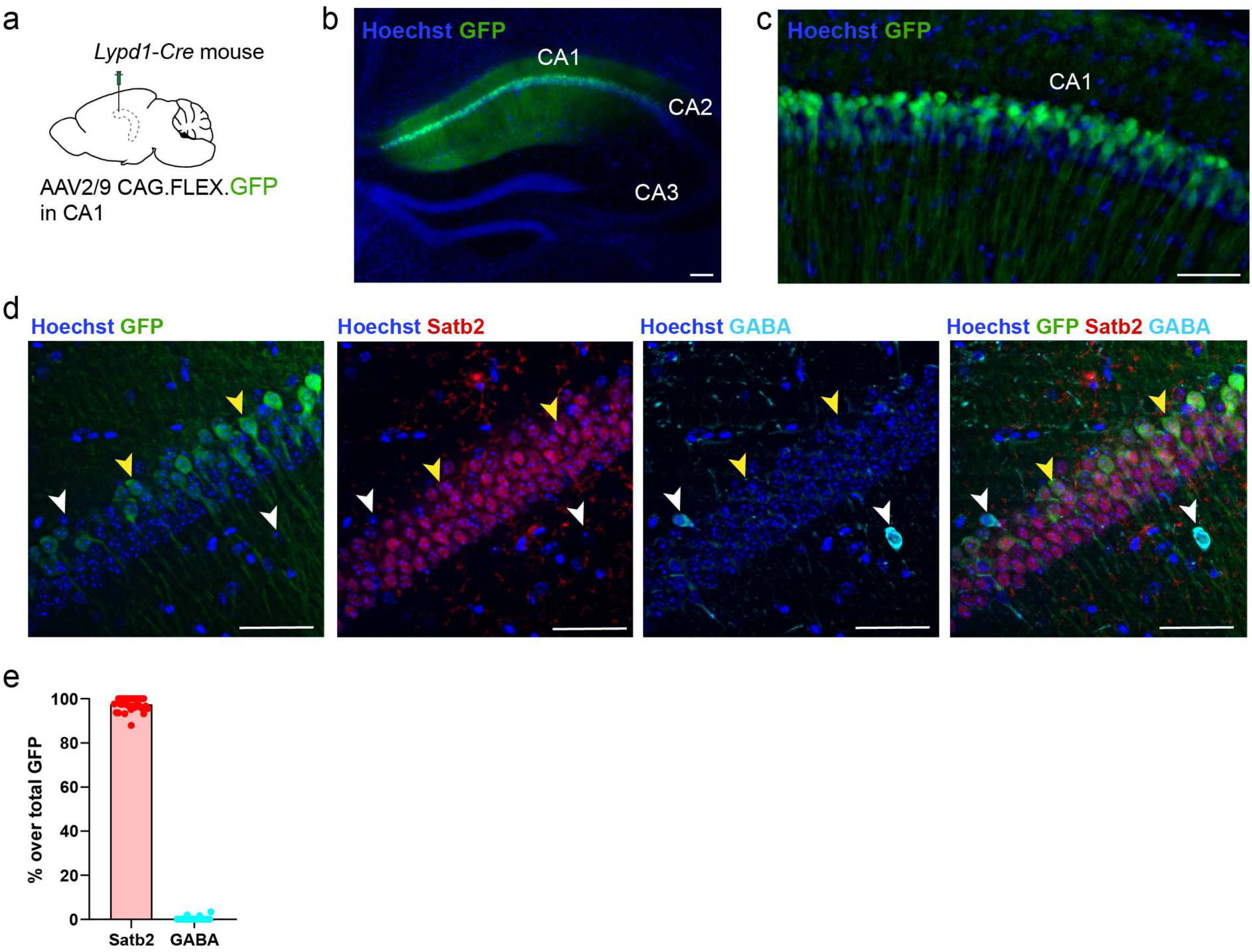
Cre expression in *Lypd1-Cre* mice is restricted to CA1 pyramidal neurons. **a.** *Lypd1-Cre* mice injected in dCA1 with AAV2/9 CAG.FLEX.eGFP.WPRE. **b-c.** Coronal section labelled for GFP and Hoechst in CA1. **d**. Immunohistochemistry against Satb2 (excitatory neurons) or GABA (interneurons) in CA1 of Lypd1-Cre mice injected with AAV2/9 CAG.FLEX.eGFP in CA1. Yellow arrowheads indicate GFP^+^ expressing Satb2. White arrowheads indicate cells positive for GABA and negative for Satb2 and GFP. **e.** Quantification of the percentage of Satb2 or GABA cells over total GFP cells in CA1. 3 mice, 400 cells/mice. Scale bars 100 µm (b) and 50 µm (c,d).

**Supplementary figure 2 related to figure 1.**
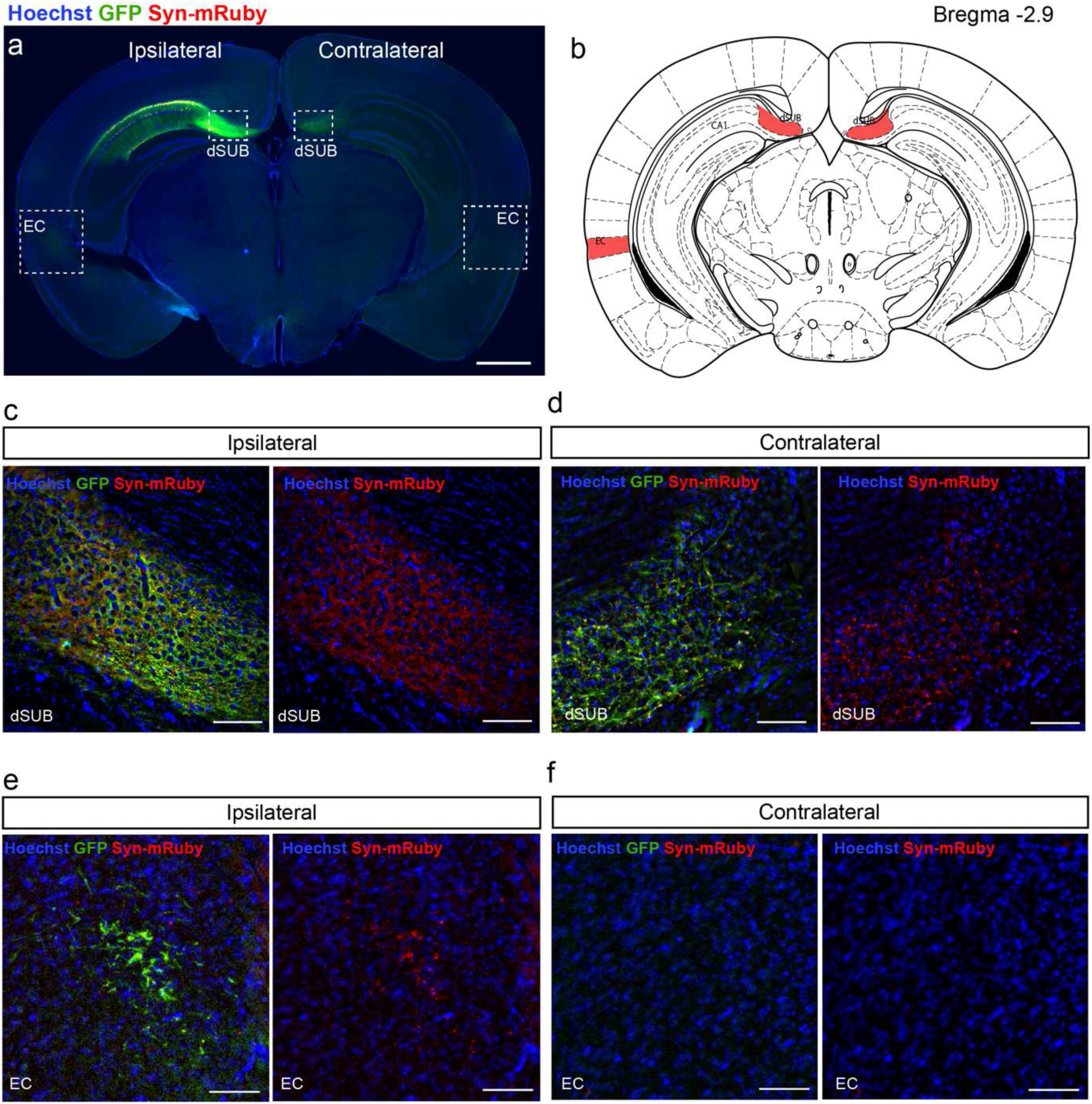
Cortical contralateral targets of dCA1 neurons. **a.** Coronal section of a *Lypd1-Cre* mouse injected in dCA1 with AAV2/DJ hSyn.FLEX.mGFP.2A.Synatophysin-mRuby and labeled for GFP and mRuby. **b.** Drawing at the level of the section shown in (a) reproduced from Paxinos et al. In red, regions in which dCA1 neurons project at this antero-posterior level. **c-f.** Magnifications of (a) showing the ipsilateral dorsal subiculum (c), contralateral dorsal subiculum (d), ipsilateral entorhinal cortex (EC, e) and contralateral entorhinal cortex (f). This experiment was reproduced in 6 mice. Scale bars: 1mm (a) and 100 µm (c-f).

**Supplementary figure 3 related to figure 1.**
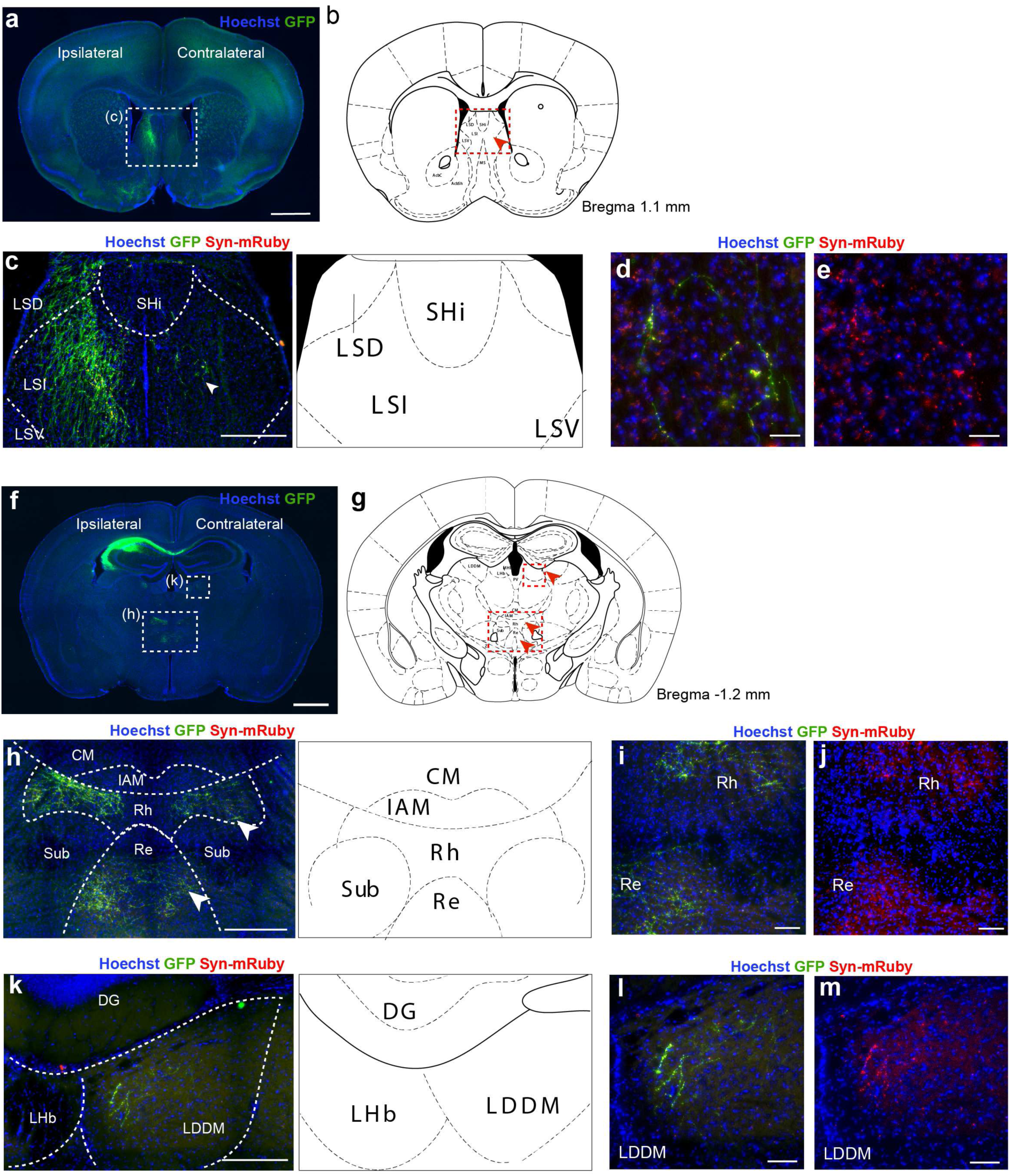
Subcortical contralateral targets of dCA1 neurons. **a.** Coronal section of a *Lypd1-Cre* mice injected in dCA1 with AAV2/DJ hSyn.FLEX.mGFP.2A.Synatophysin-mRuby and labelled for GFP and mRuby. **b.** Drawing reproduced from Paxinos et al. at the level of the section shown in (a). The red arrowhead indicates the contralateral target of dCA1 at this level. **c-e.** Magnification from (a). The white arrowhead shows terminals in contralateral rLS. **d-e.** Magnification from (c) showing contralateral rdLS. **f.** Coronal section showing the injection site and thalamic targets of dCA1. **b.** Drawing reproduced from Paxinos et al. at the level of the section shown in (f). The red arrowhead indicates the contralateral target of dCA1 at this level. **h.** Magnification from (f). The white arrowheads show terminals in contralateral Rh and Re nuclei. **i-j.** Magnification from (h) showing contralateral Rh and Re nuclei. **k.** Magnification from (f) showing the contralateral LDDM. **l-m.** Magnifications from (k) showing fibers and terminals in contralateral LDDM. This experiment was reproduced in 6 mice. Scale bars: 1mm (a,f), 300 µm (c,h), 200 µm (k), 100 µm (i,j) and 50 µm (d,e,l,m). (LSD) dorsal lateral septum, (SHi) septohippocampal nucleus, (LSI) intermediate lateral septum, (LSV) ventral lateral septum, (CM) central medial nucleus of the thalamus, (Sub) submedial nucleus of the thalamus, (Re) nucleus of reuniens, (Rh) rhomboid nucleus, (IAM) inter-mediodorsal nucleus of the thalamus, (LDDM) laterodorsal nucleus of the thalamus, (LHb) lateral habenula and (DG) dentate gyrus.

**Supplementary figure 4 related to figure 1.**
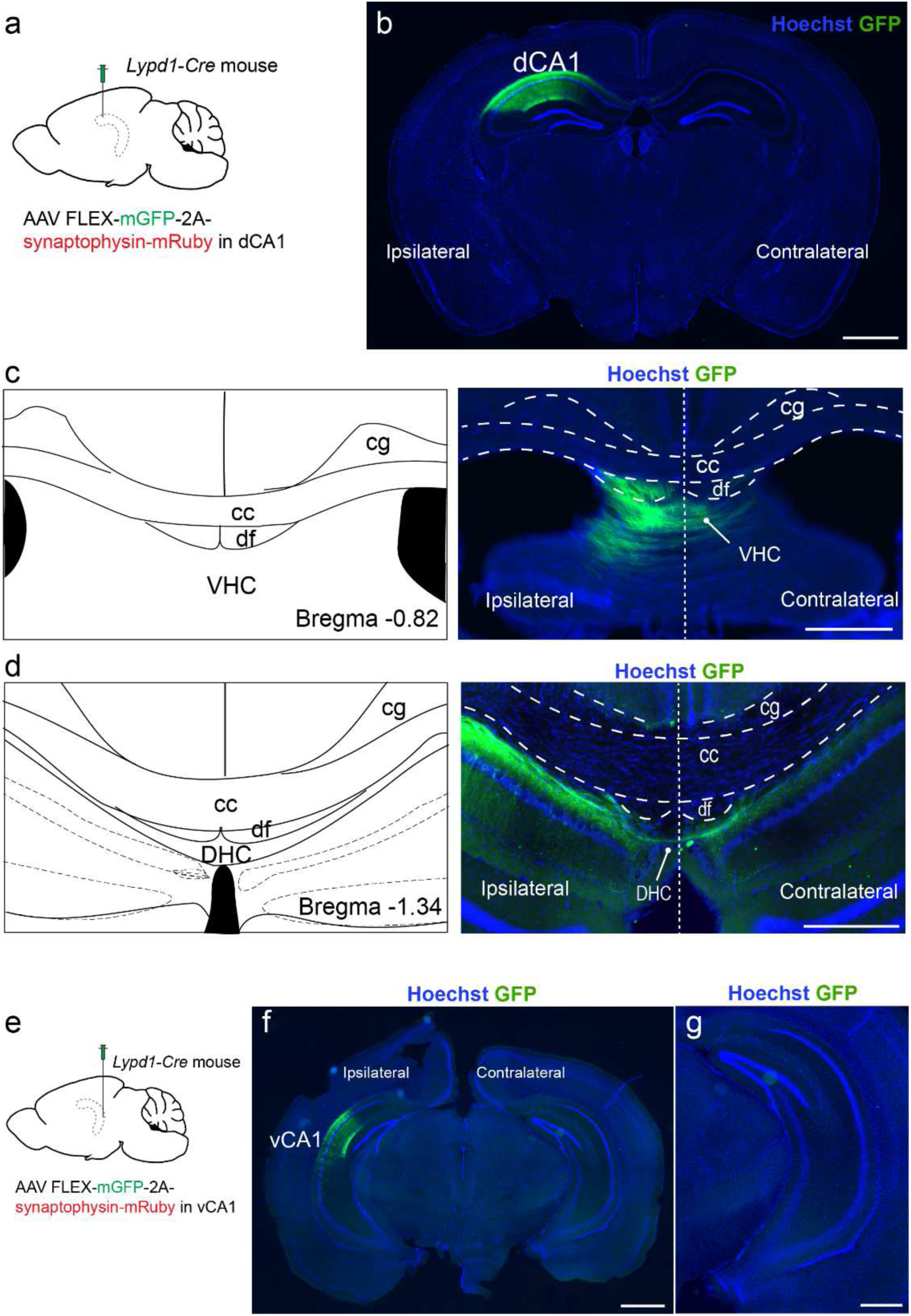
Hippocampal contralateral projections from dCA1. **a.** *Lypd1-Cre* mice injected in dCA1 with AAV2/DJ hSyn.FLEX.mGFP.2A.Synatophysin-mRuby. **b.** Coronal section labelled for GFP. **c.** Coronal section from the same brain showing the VHC. **d.** Coronal section from the same brain showing the dorsal hippocampal commissures. **e.** *Lypd1-Cre* mice injected in ventral CA1 (vCA1) with AAV2/DJ hSyn.FLEX.mGFP.2A.Synatophysin-mRuby. **f.** Coronal section labeled for GFP. **g.** Magnification of the contralateral hemisphere shown in (f). This experiment was reproduced in 6 mice. Scale bars: 1mm (b,f) and 500 µm (c,d,g). (cg) cingulum bundle, (cc) corpus callosum, (df) dorsal fornix, (DHC) dorsal hippocampal commissure and (VHC) ventral hippocampal commissure.

**Supplementary figure 5 related to figure 1.**
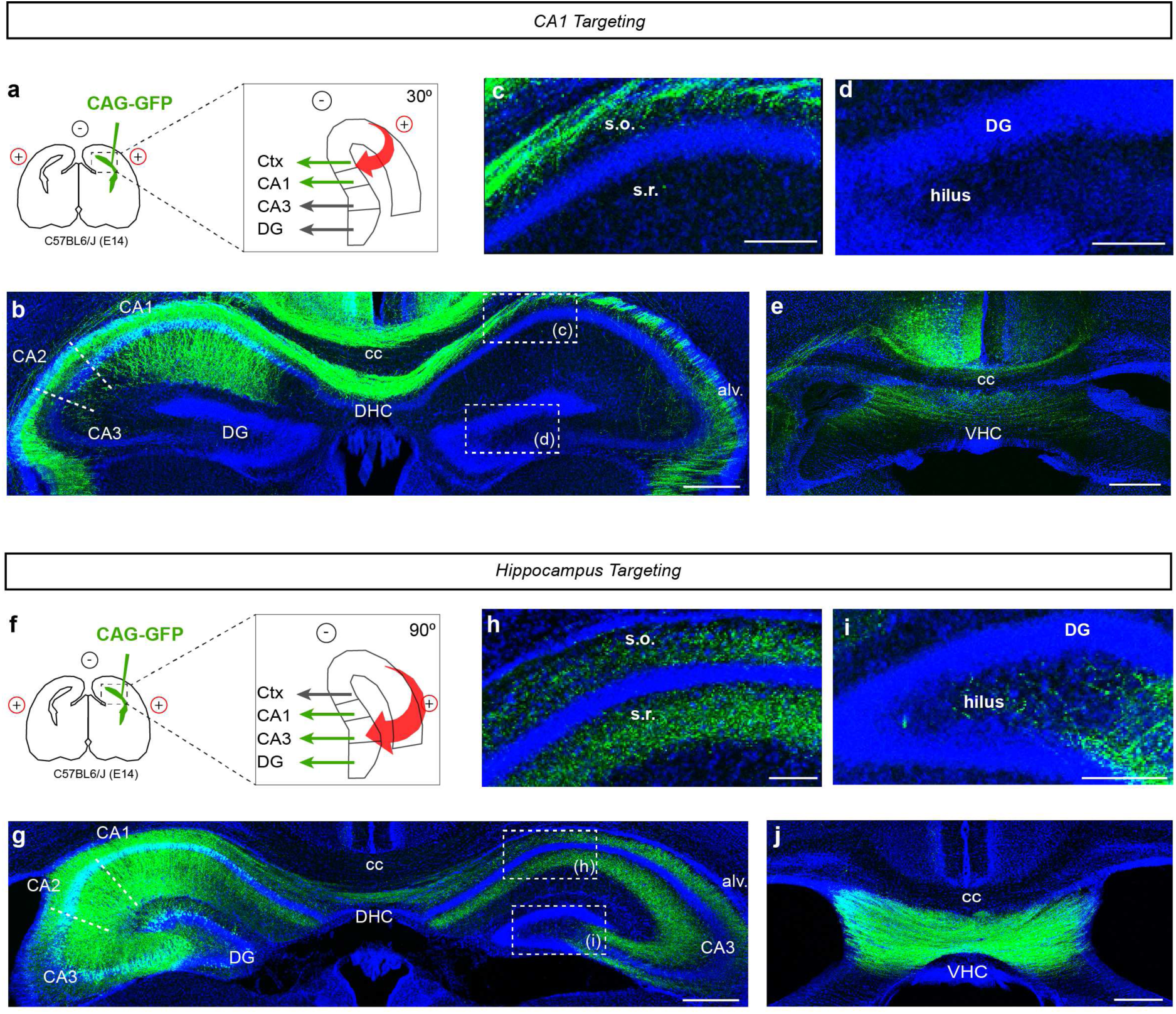
Development of hippocampal interhemispheric projections. **a.** Schematic of in utero electroporation (IUE) of WT mice at E14 with the third electrode system to specifically target CA1 pyramidal neurons in the hippocampus. **b.** Coronal section of the hippocampus at P7 following IUE as in (a) labelled for GFP. **c-d.** Magnification of the contralateral CA1 (c) and DG (d). **e.** Coronal section of the VHC of the same brain. **f.** Schematic of IUE of WT mice at E14 with the third electrode system to target pyramidal neurons of the entire hippocampus. **g.** Coronal section of the hippocampus at P7 following IUE as in (f) labelled for GFP. **h-i.** Magnifications of the hilus of the contralateral CA1 (f) and DG (i). **j.** Coronal section of the VHC of the same brain. Scale bars: 400 µm (b,e,g,j) and 100 µm (c, d, h, i).

**Supplementary figure 6 related to figure 2:**
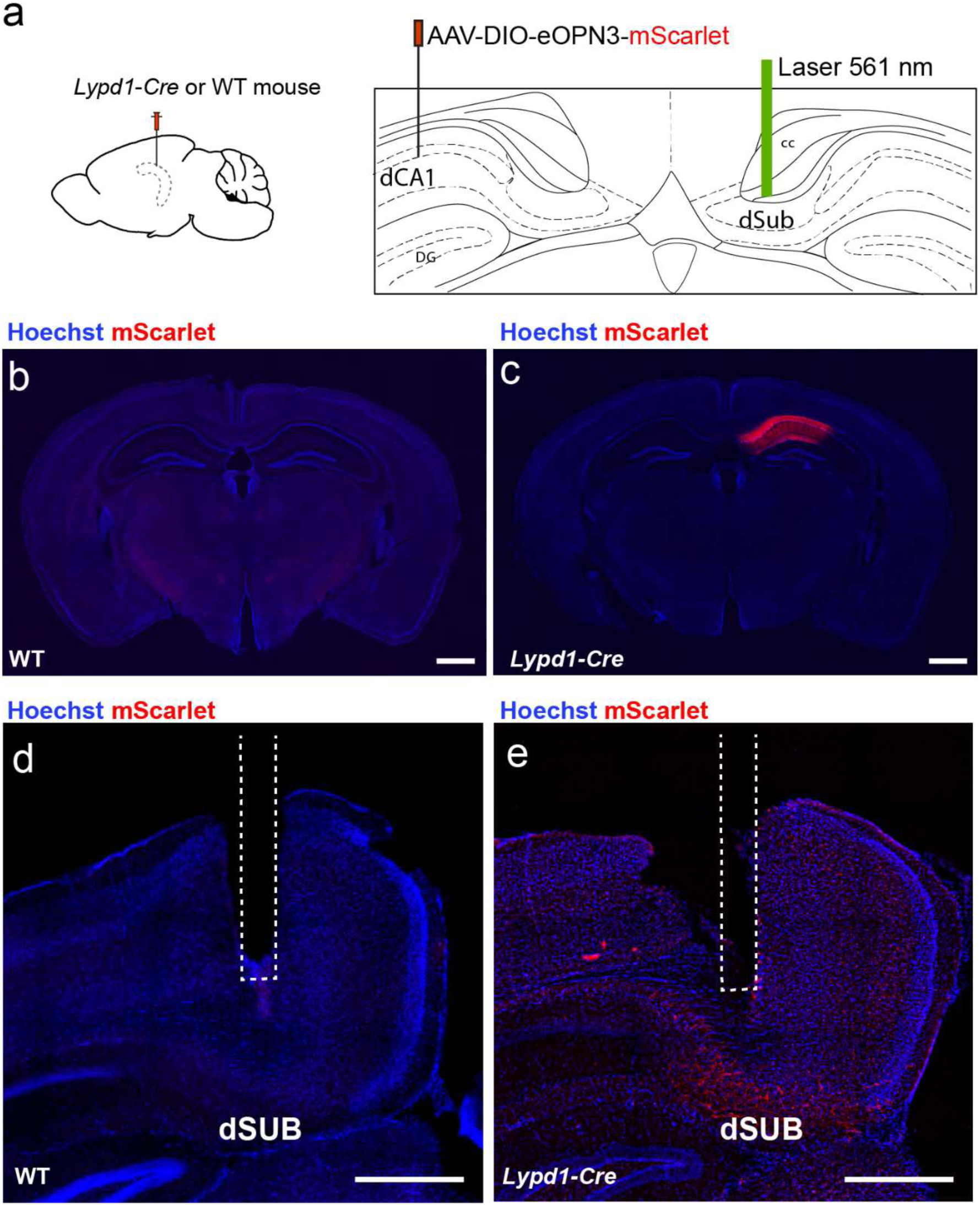
eOPN3 expression and implants. **a.** *Lypd1-Cre* and WT mice from both sexes injected in the right dCA1 with AAV2/1 hSyn1-DIO-eOPN3-mScarlet-WPRE and implanted with an optic fiber above the left dSUB to silence dCA1 to dSUB interhemispheric terminals. **b-c.** Coronal section of WT (b) or *Lypd1-Cre* (c) mouse brain labelled for mScarlet. **d-e.** Coronal section of WT (d) or *Lypd1-Cre* (e) mouse showing the lens implant above the left dSUB of the same brains shown previously. Scale bars: 1mm (b-c), 500 µm (d-e).

**Supplementary figure 7 related to figure 2:**
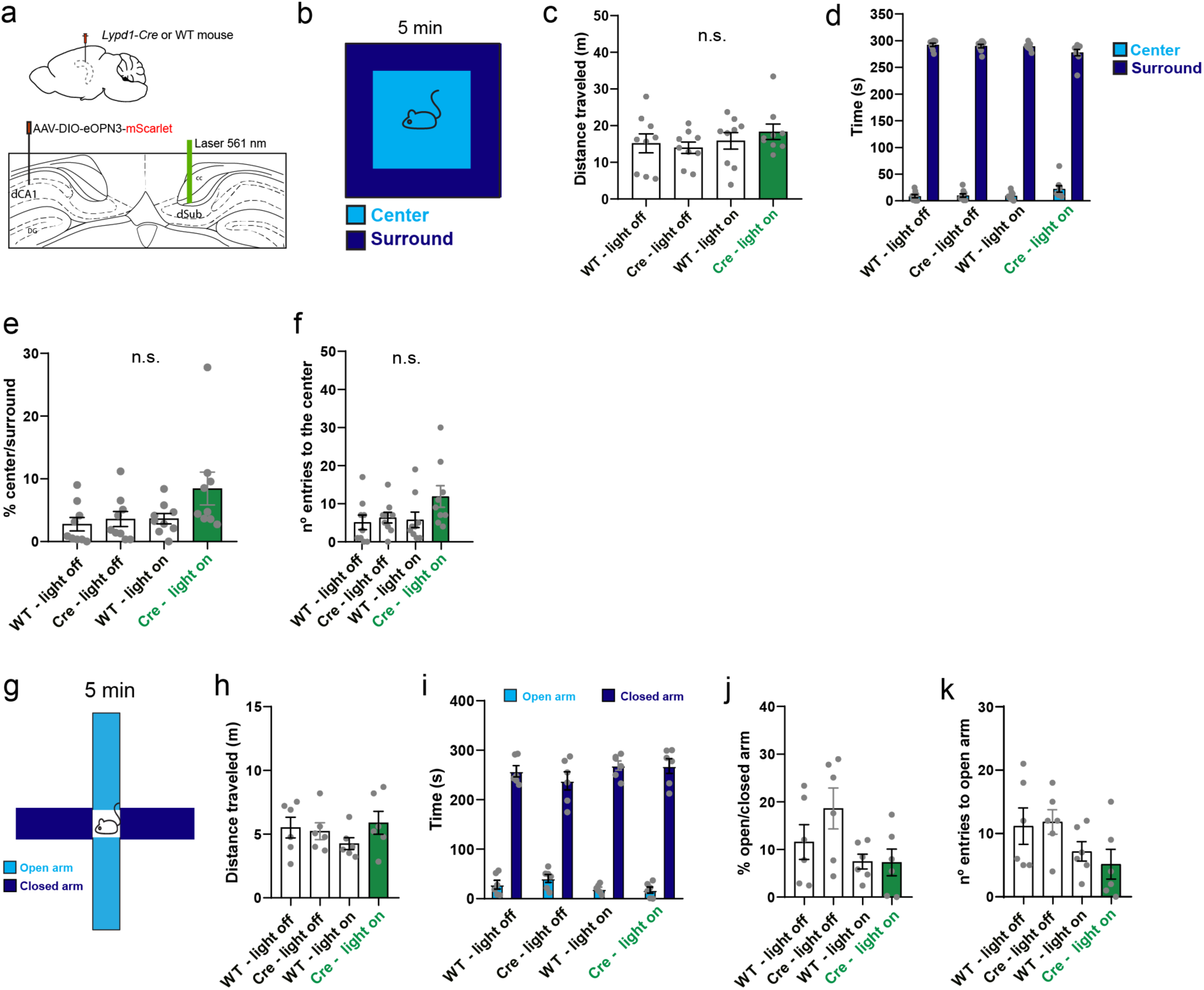
Open field and elevated plus-maze tests. **a.** *Lypd1-Cre* and WT mice from both sexes injected in the right dCA1 with AAV2/1 hSyn1-DIO-eOPN3-mScarlet-WPRE and implanted with an optic fiber above the left dSUB to silence dCA1 to dSUB interhemispheric terminals. **b.** Schematic of the open field test. **c.** Total distance traveled during the open field test. For (c-f), each point corresponds to one mouse (9 WT and 9 *Lypd1-Cre* mice). **d.** Time spent in the center or surround of the open field. **e.** Ratio of the time spent in the center/surround. **f.** Number of entries into the center zone in each group. **g**. Schematic of the elevated plus maze test (EPM). **h.** Total distance traveled during the EPM. For (h-k), each point corresponds to one mouse (6 WT and 6 *Lypd1-Cre* mice). **i.** Time spent in the open or closed arms during the EPM. **j.** Ratio of the time spent in the open/closed arms. **k.** Number of entries into the open arm in each group. For the entire figure, bar graphs represent mean ± SEM..

**Supplementary figure 8 related to figure 2:**
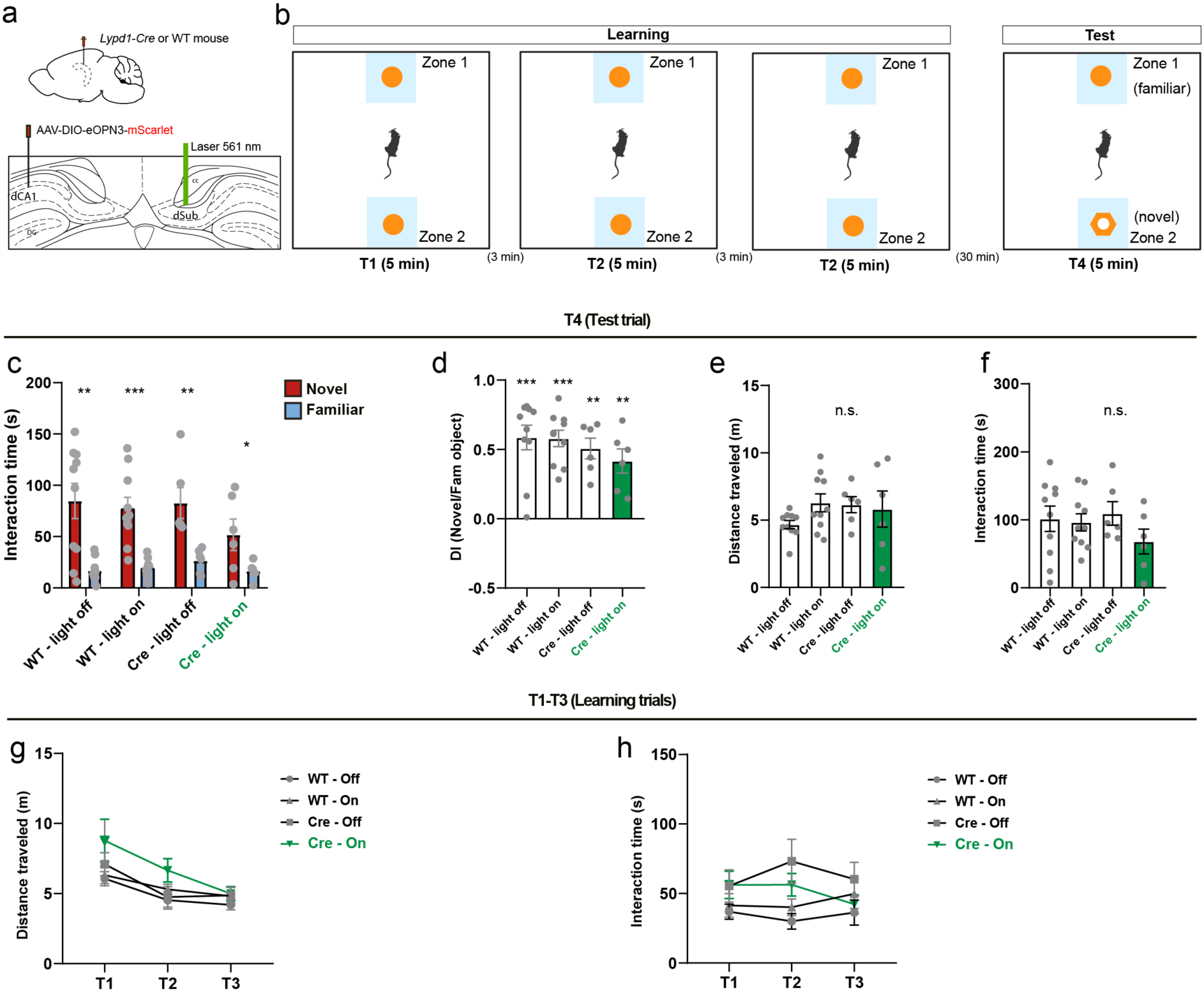
Novel object recognition test. **a.** Schematic of the experiment. *Lypd1-Cre* or WT mice from both sexes injected in the right dCA1 with AAV2/1 hSyn1-DIO-eOPN3-mScarlet-WPRE and implanted with an optic fiber above the left dSUB to silence dCA1 to dSUB interhemispheric terminals. **b.** Schematic of the novel object recognition test. **c.** Time of interaction with the familiar or novel object. For (c-f), each point corresponds to one mouse (10 WT and 6 *Lypd1-Cre* mice). **d.** Discrimination index of the novel over the familiar object. **e.** Distance traveled during the test trial (T4). **f.** Interaction time with the object during the test trial (T4). **f.** Distance traveled during the learning trials (T1-T3). **h.** Interaction time with the object during the learning trials (T1-T3). For the entire figure, bar graphs represent mean ± SEM.

**Supplementary figure 9 related to figure 4:**
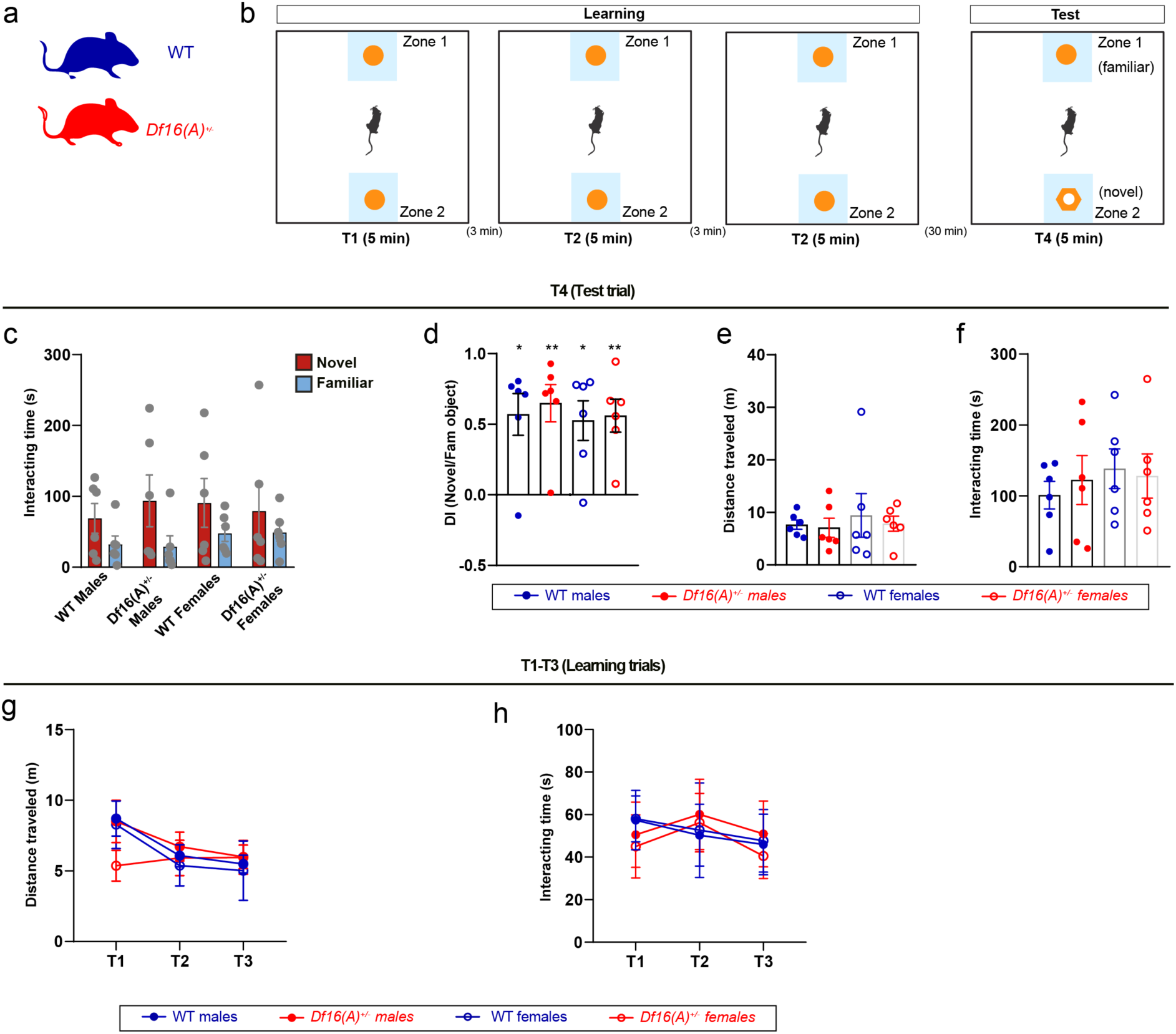
Novel object recognition test female and male *Df16(A)^+/-^* mice. **a.** Female and male *Df16(A)^+/-^* or WT mice were tested. **b.** Schematic of the novel object recognition test. **c.** Time of interaction with the familiar or novel object. For (c-f), each point corresponds to one mouse (6 mice per group). **d.** Discrimination index of the novel over the familiar object. **e.** Distance traveled during the test trial (T4). **f.** Interaction time with the object during the test trial (T4). **g.** Distance traveled during the learning trials (T1-T3). **h.** Interaction time with the object during the learning trials (T1-T3). For the entire figure, bar graphs represent mean ± SEM.

**Supplementary figure 10 related to figures 5 & 6:**
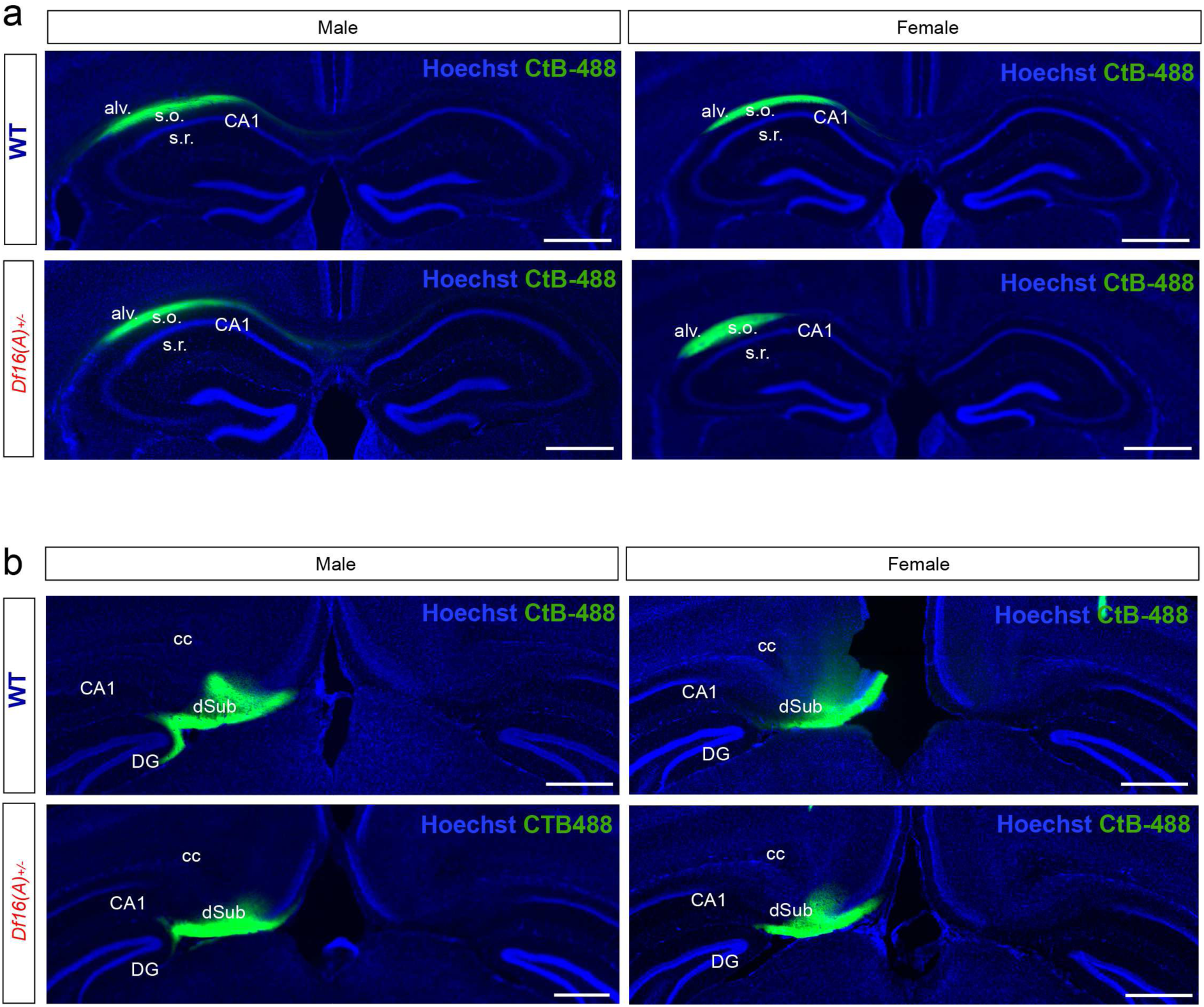
CtB-488 injection sites. **a.** Coronal sections showing representative CtB-488 injections in dCA1 of the right hemisphere in male or female WT or *Df16(A)^+/-^* mice. **b.** Coronal sections showing representative CtB-488 injections in dSUB of the right hemisphere in male or female WT or *Df16(A)^+/-^* mice. Scale bars: 500 µm.

**Supplementary figure 11 related to figure 5:**
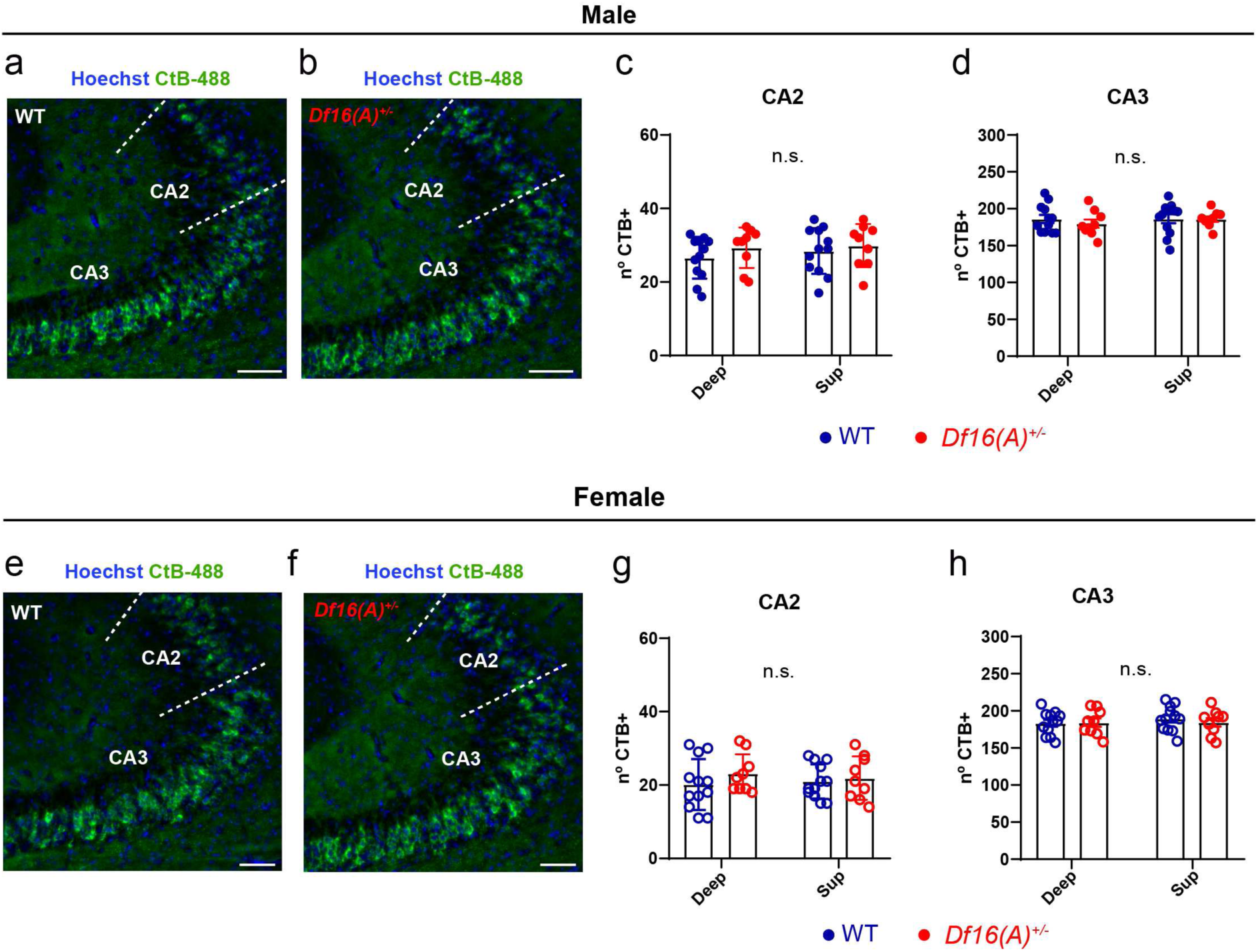
CtB^+^ cells in contralateral hippocampus. a-b. CtB^+^ cells in the left hippocampus of WT and *Df16(A)^+/-^* male mice after CtB-488 injections in the right dCA1. **c.** Number of CtB^+^ cells in deep and superficial layers of CA2. **d.** Number of CtB^+^ cells in deep and superficial layers of CA3. For (c-d), each point represents one observation (4 WT and 3 *Df16(A)^+/-^* mice, 3 observation per mouse). **e-f.** CtB^+^ cells in the left hippocampus of WT and *Df16(A)^+/-^* female mice following injection in right dCA1. **g.** Number of CtB^+^ cells in deep and superficial layers of CA2. **h.** Total number of CtB^+^ cells in deep and superficial layers of CA3. For (c-d), each point represents one observation (4 WT and 3 *Df16(A)^+/-^* mice, 3 observation per mouse). For the entire figure, bar graphs represent mean ± SEM.

**Supplementary figure 12 related to figures 5 and 6:**
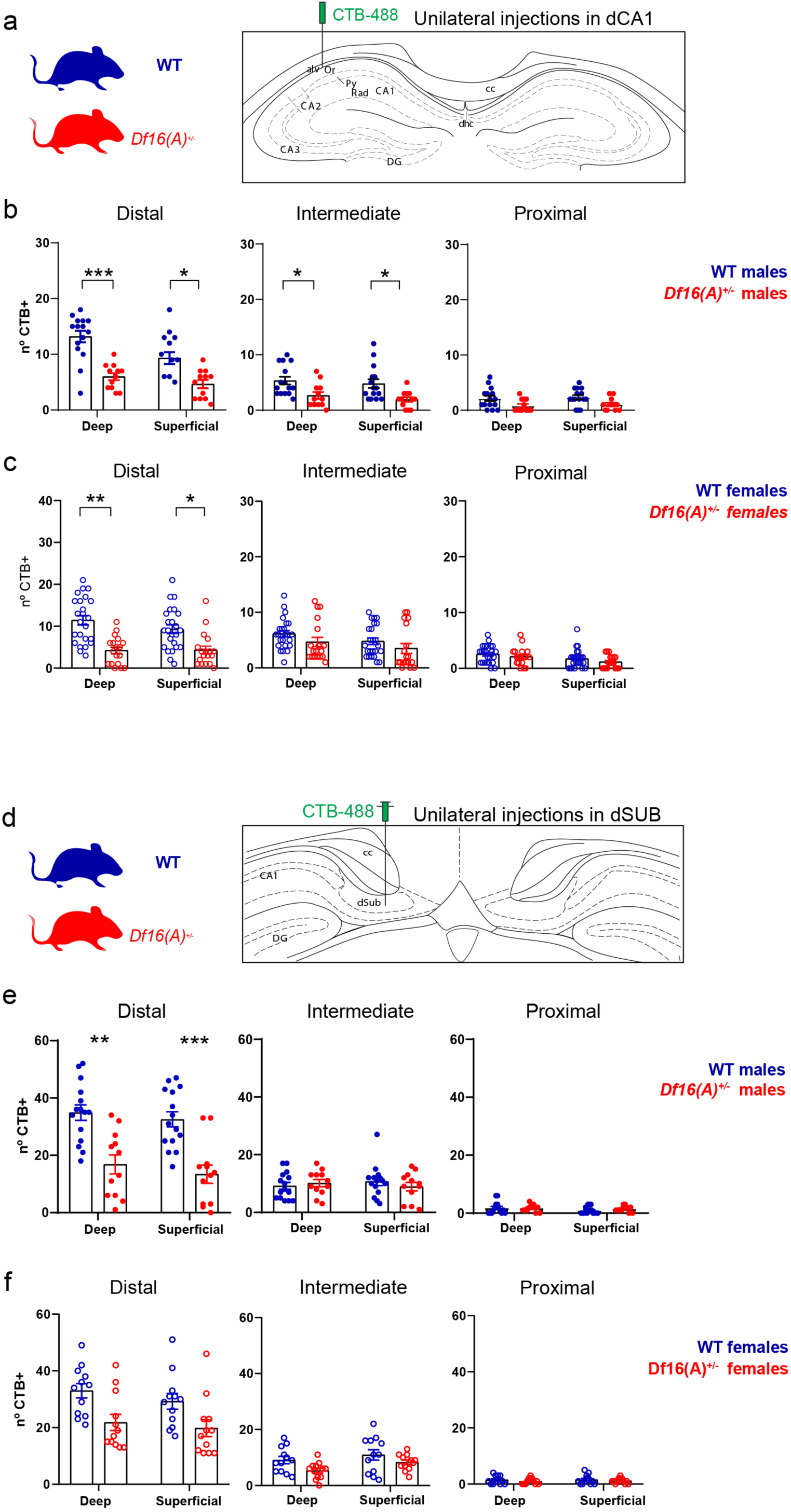
CtB^+^ cells in deep and superficial layers of contralateral CA1 after dCA1 or dSUB injections. **a.** WT and *Df16(A)^+/-^* mice injected with CtB-488 in the right dCA1. **b.** Number of CtB^+^ cells in deep and superficial layers of proximal, intermediate and distal contralateral dCA1 of WT and *Df16(A)^+/-^* male mice. Each point represents one observation (5 WT and 4 *Df16(A)^+/-^* mice, 3 observations per mouse). **c.** Number of CtB^+^ deep and superficial layers of proximal, intermediate and distal contralateral dCA1 of WT and *Df16(A)^+/-^* female mice. Each point represents one observation (8 WT and 6 *Df16(A)^+/-^* mice, 3 observations per mouse). **d.** WT and *Df16(A)^+/-^* mice injected with CtB-488 in the right dSUB. **e.** Number of CtB^+^ cells in deep and superficial layers of distal, intermediate and proximal contralateral dCA1 of WT and *Df16(A)^+/-^* male mice. Each point represents one observation (5 WT mice and 4 *Df16(A)^+/-^* mice, 3 observations per mouse). **f.** Number of CtB^+^ cells in deep and superficial layers of distal, intermediate and proximal contralateral dCA1 of WT and *Df16(A)^+/-^* male mice. Each point represents one observation (4WT and 4 *Df16(A)**^+/-^*** mice, 3 observations per mouse). For the entire figure, bar graphs represent mean ± SEM.

